# Membrane cliffs are giant, recursive platforms that drive calcium and protein kinase signaling for cell growth

**DOI:** 10.1101/2024.01.08.574622

**Authors:** Marco Trerotola, Valeria Relli, Romina Tripaldi, Pasquale Simeone, Emanuela Guerra, Andrea Sacchetti, Martina Ceci, Ludovica Pantalone, Paolo Ciufici, Antonino Moschella, Valeria R. Caiolfa, Moreno Zamai, Saverio Alberti

## Abstract

The transmembrane glycoproteins Trop-1/EpCAM and Trop-2 independently trigger Ca^2+^ and kinase signals for cell growth and tumor progression. We discovered that Trop-1 and Trop-2 are recruited at overlapping sites at free cell edges. Z-stack analysis and three-dimensional reconstruction of these sites revealed previously unrecognized, protruding membrane regions ≥20 µm-long, up to 1.5 µm high, then named ‘cliffs’. Cliffs appeared confined to essentially immobile sites of the cell membrane, where they recursively assembled over hundreds of seconds. Cliffs were shown to recruit growth-driving kinases and downstream cytoplasmic effectors. Trop-2 stimulates cell growth through a membrane super-complex that comprises CD9 and PKCα. Our findings indicated that the growth-driving Trop-2 super-complex assembles at cliffs. Cliffs acted as sites of phosphorylation/activation of growth-driving kinases and as origins of Ca^2+^ signaling waves, indicating cliffs as novel signaling platforms for drivers of cell growth. Cliffs were induced by growth factors and disappeared upon growth factor deprivation, suggesting cliffs as pivotal platforms for signaling for cell growth.

## Introduction

The transmembrane glycoproteins Trop-1/EpCAM and Trop-2 have been shown to drive tumor growth ^1–3^ and metastatic diffusion ^4,5^. Trops drive tumor progression upon overexpression as wild-type molecules ^1,2,6,7^, which are activated by ADAM10 cleavage at a conserved position in the thyroglobulin domain ^4,8–11^. Trop proteolytic cleavage triggers a downstream proteolytic cascade, carried out by the TNF-α-converting enzyme (TACE) followed by γ-secretase cleavage within the transmembrane domain, which leads to nuclear signaling and transcription factor activation ^4,10,12,13^. Trop-2 was shown to activate a ubiquitous set of signaling molecules, that subsequently modulate the basal growth programs of cancer cells. This protein super-complex is assembled around CD9, Trop-2 ^14^ and the Na+/K+-ATPase ion pump. Activation of the Trop-2 supercomplex leads to the release of intracellular Ca^2+^ ^14,15^ and to the recruitment of PKCα to the cell membrane, for phosphorylation of the Trop-2 cytoplasmic tail. This establishes a feed-forward activation loop with remodeling of the β-actin/α-actinin/myosin II cytoskeleton, through cofilin-1, annexins A1/A6/A11 and gelsolin. This drives malignant progression through the cleavage of the β-actin-binding site of E-cadherin ^14^, and downstream activation of Akt ^16^, ERK, NFkB, and cyclin D1 ^17^.

We discovered that the Trop super-complex is assembled at newly identified, protruding membrane regions, ≥20 µm long and up to 1.5 µm high, which were named cliffs. Cliffs recursively assembled at conserved, essentially immobile sites of the cell border and recruited growth-driving kinases and downstream cytoplasmic effectors. Cliffs were shown to operate as signaling platforms for drivers of cell growth, through kinase phosphorylation/activation and as trigger sites for waves of Ca^2+^ signals. Cliffs were shown to be induced by growth factors and to wane upon growth factor deprivation, suggesting cliffs to play a pivotal role in signaling for cell growth.

## Results and discussion

### Cell growth drivers localize at overlapping sites of the cell membrane

Trop-1/EpCAM and Trop-2 were previously shown to trigger Ca^2+^ signals and to drive cancer cell growth ^1,2,5,6,8,14,15^. We noticed that Trop-1 and Trop-2, despite distinct intra-cellular distribution patterns (Figure S1), were recruited at overlapping sites at the cell membrane (Figure 1). This finding was far from trivial, as Trop-1 is not required for Trop-2 signaling ^5,7,14^, first suggesting a broader functional relevance of these co-recruitment sites.

**Figure 1.**
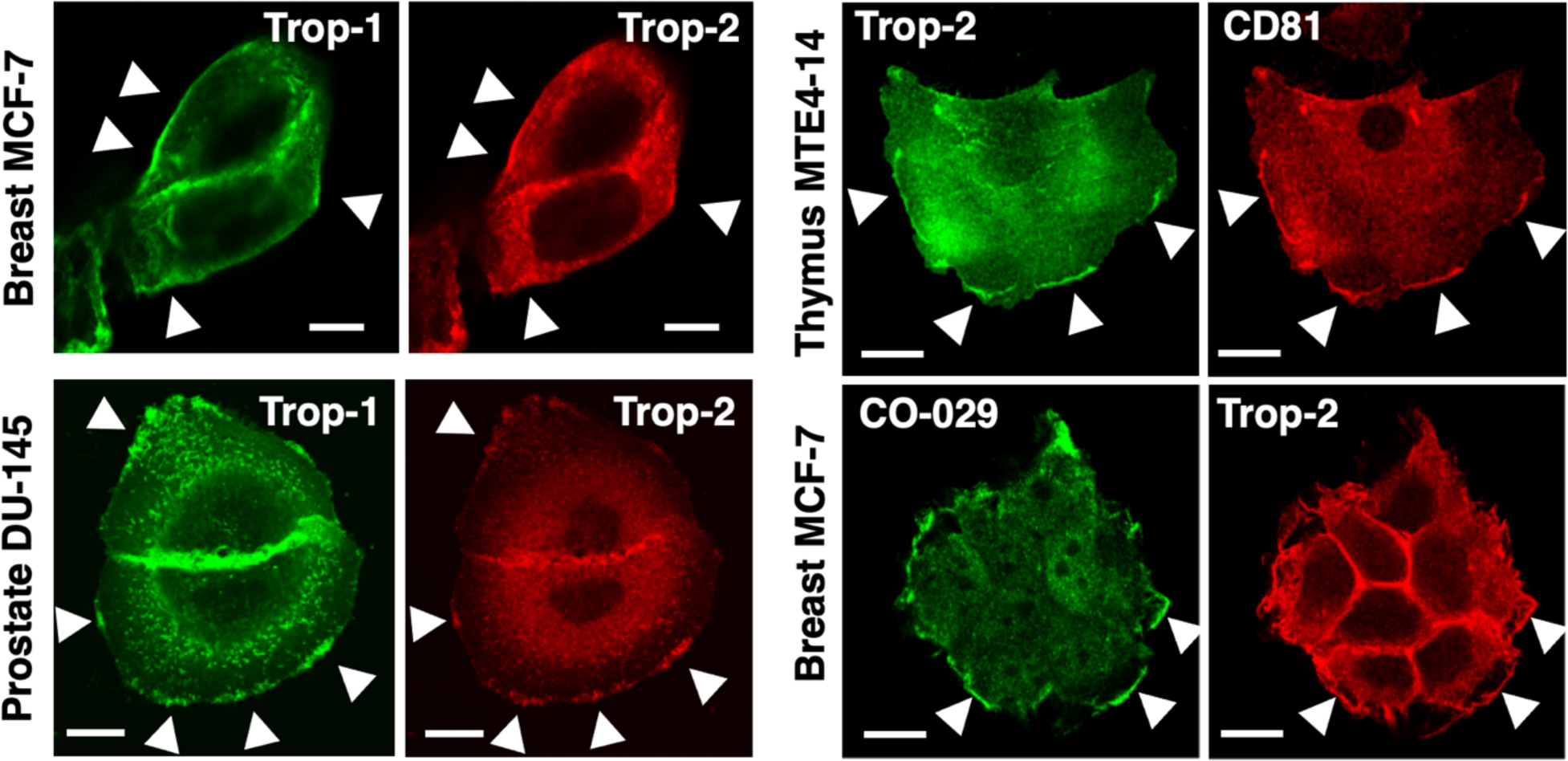
Trop-1, Trop-2, CD81, CO-029 are co-recruited at distinct membrane segments. Breast MCF-7, prostate DU-145 cancer cells and transformed MTE4-14 thymus cells were analyzed for expression and localization of signal transducers by confocal microscopy. White arrowheads indicate colocalization at distinct membrane segments. (*left*) Trop-1 and Trop-2 were revealed with the HT29/26-Alexa488 and T16-Alexa633 mAb, respectively. (*right*) Colocalization analysis of Trop-2 with the CD81 and CO-029 tetraspanins.

Recruitment of Trop-1 and Trop-2 at the cell membrane is essential for functional activation ^4,5,10,11,13,18^, through the assembly of a signaling super-complex ^14^. Signaling platforms function as membrane sites that colocalize molecular signals, thus facilitating their interaction ^19,20^. We thus explored whether Trop recruitment sites of the cell membrane operated as signaling platforms for the Trop super-complex. Using FP chimeras for parallel imaging and functional analysis, we showed that Trop-1-FP and Trop-2-FP were efficiently transported to the cell membrane, at sharply confined sites (Figure 2A-C). Care was taken not to overexpress FP-chimera, but to confine expression to average expression levels in tumor cells ^14^. Trop-1–FP and Trop-2–FP were shown to stimulate cell growth as efficiently as comparable levels of the wild-type molecules (Figure 2A), indicating functional competence for cell signaling. We went on to assess whether other key components of the Trop signaling super-complex were co-recruited at the same membrane sites. Non-parametric Spearman correlation analysis of fractional perimeter segments occupied by any one molecule at any specific time was performed (Table S1). Trop-1-FP, Trop-2-FP, CD9-FP and ERK-FP were found to tightly colocalize at the cell membrane (Movies S1, S2), with extremely high correlation coefficients (0.986-0.993; P <0.0001) (Table S1), supporting the notion that these co-recruitment sites may operate as functional platforms for signaling. Additional classes of signaling protein-FP chimeras appeared recruited at these same membrane regions. Recruited molecules included actin-bundling determinants (fascin-GFP), cytoplasmic kinases (PKCα-GFP) and tetraspanins (CD316-mCherry) (Figure 2). On the other hand, PKCδ-GFP (Figure 2C) and PKCε-GFP were not transported to PKCα-GFP membrane recruitment sites, suggesting co-recruitment specificity.

**Figure 2.**
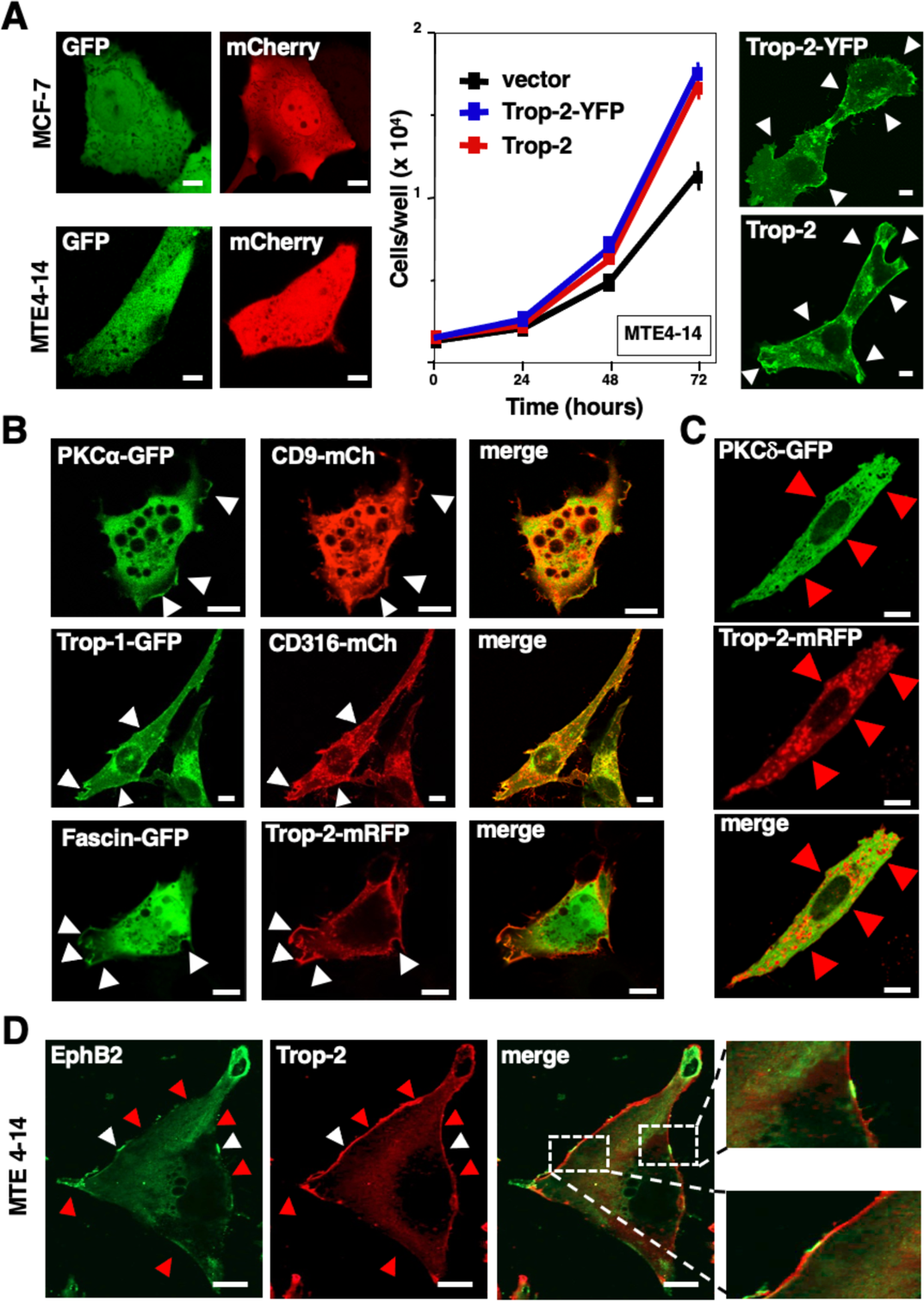
Distinct signaling molecules are co-recruited at confined cell membrane regions. Thymus MTE4-14 and breast MCF-7, MDA-MB-231 cells were transfected with signaling molecule-FP chimeras or Ca^2+^ indicators. Membrane recruitment and impact on cell growth were assessed. White arrowheads indicate colocalization of the signal transducers that were challenged. Red arrowheads indicate absence thereof. (**A**) (*left*) Control transfectants show absence of mCherry or GFP tag localization to the cell membrane in MCF-7 and MTE4-14 cells. (*mid*) The Trop-2–YFP chimera was shown to induce cell growth just as well as wild-type Trop-2. Data are presented as mean±SEM. (*right*) Parallel localization of Trop-2–YFP chimeras versus wild-type Trop-2, as revealed by T16 anti-Trop-2 mAb-staining in MTE4-14 transfectants. (**B**) Horizontal panel strips show membrane colocalization of PKCα–GFP and CD9–mCherry, or Trop-1–GFP and CD316–mCherry, or Fascin–GFP and Trop-2– mRFP in MTE4-14 transfectants. (**C**) Lack of colocalization of Trop-2–mRFP versus PKCδ–GFP in MTE4-14 cells. Scale bars, 10 µm. (**D**) Absence of colocalization of Trop-2 with EphB2 at cliffs. ROI magnify details of different membrane localization regions. Bars, 10 µm.

Analysis of endogenous molecules provided corresponding findings to those obtained with FP chimeras. Utilizing Trop-2 and CD9 as membrane-recruitment-site tracers, lack of co-recruitment was shown for EphB2 (Figure 2D), CD34, E-cadherin among transmembrane molecules, caveolin-1, β-catenin among cytoplasmic molecules, thus supporting a model of selective co-recruitment of specific signal transducers at these sites. Among tetraspanins, we observed tight colocalization and co-capping of Trop-2 with CD9, CO-029, CD81 and CD98. On the other hand, limited colocalization was detected with CD316 and essentially none was found with CD151 (Figure 2B), suggesting a fine regulation of the recruitment of different classes of tetraspanins at these membrane sites ^14^.

### The cliff membrane regions

Z-stack analysis and three-dimensional (3D) reconstruction of the membrane Trop-1, Trop-2, CD9 recruitment sites were used to explore the structure of the corresponding membrane regions (Movie S3). This also allowed us to prevent artifacts due to trivial shifts of motile membranes across neighbouring confocal sections. The 3D analyses revealed previously unrecognized, protruding membrane regions (up to 1.5 µm high) (Figure 3A) at conserved sites of the cell border. These membrane regions were thus named ‘cliffs’ (Box 1).

**Figure 3.**
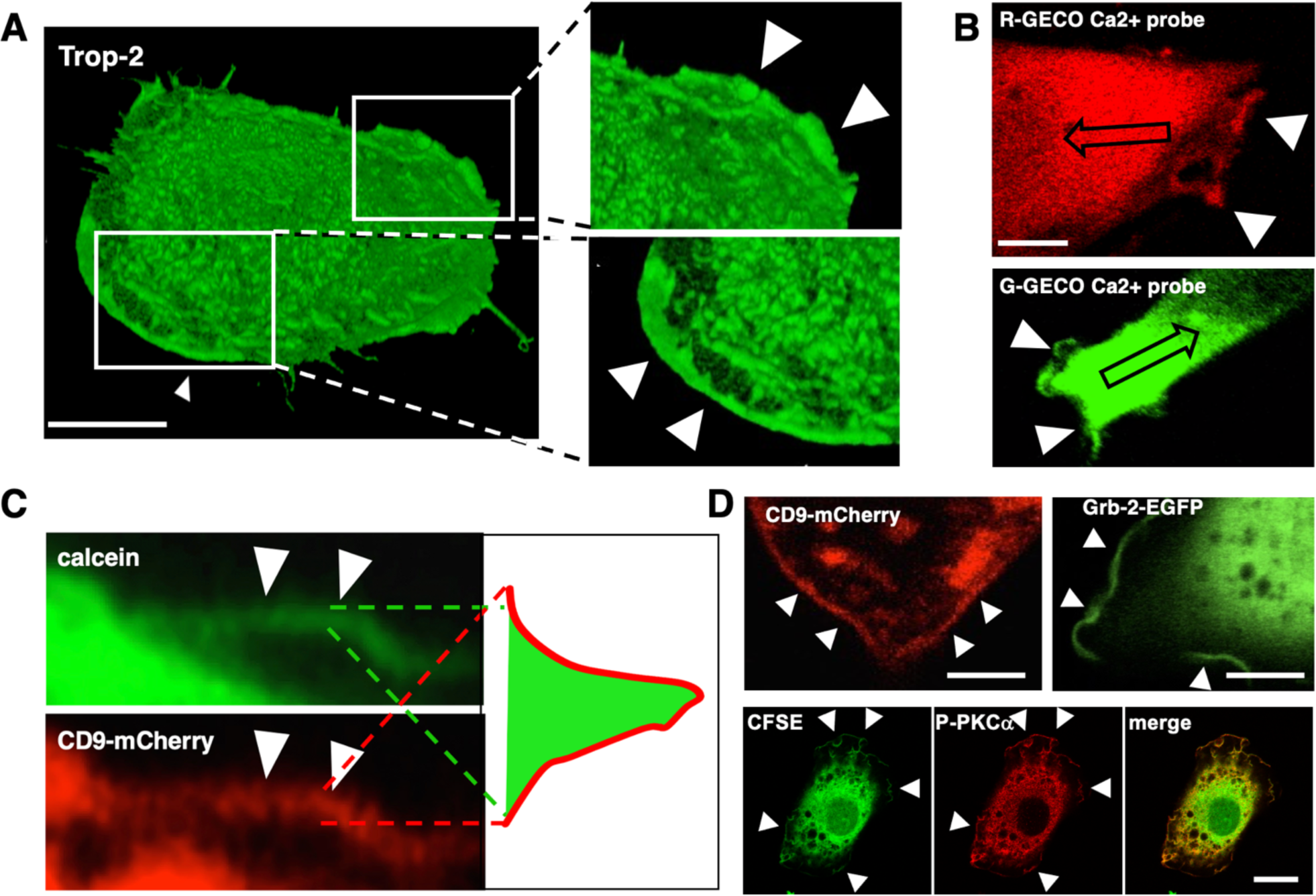
3D structure of membrane cliffs. (**A**) Confocal Z-stack 3D-reconstruction of a Trop-2-expressing MTE4-14 living cell stained with the T16 anti-Trop-2 mAb (outside membrane staining). (*insets*) Magnified regions of interest (ROI, white rectangles). Membrane cliffs are indicated by arrowheads. Bar, 5 µm. The full dataset is presented in Movie S3. (**B**) MDA-MB-231 cells and MTE4-14/Trop-2 transfectants super-transfected with the Ca^2+^ indicators G-GECO and R-GECO, respectively, to visualize cell membrane cliffs. The pictures are Movie frames taken at the time of origin of cytoplasmic Ca^2+^ waves, elicited by mAb-induced Trop-2 cross-linking ^15^. The direction of propagation of the Ca^2+^ waves is indicated by the arrows. The full dataset is presented in Movie S6. Scale bars, 5 µm. (**C**) MTE4-14 cells transfected with CD9–mCherry and treated with calcein as a cytoplasm tracer. Arrowheads indicate the same cliff region in the two paired panels. Top: The rim of cytoplasm contained in the cliff is indicated by the calcein signal. Bottom: CD9-mCherry labels the external membranes of the cliff. A cliff model is depicted in the right side of the cartoon. (**D**) (*top*) MTE4-14 cells transfected with CD9–mCherry or with Grb2-EGFP. (*bottom*) The cytoplasm of MTE4-14 cells was labeled with CSFE 5 µM in PBS 15 min at 37°C (green). Endogenous P-PKCα was stained with Alexa546 mAb (red). Merged signals are indicated on the right. Membrane cliffs are indicated by arrowheads. Bars, 5 µm.

Extensive 3D reconstructions showed that cliff occurrence and localization correlated with neither the extent of cell motility nor the direction of cell movement (Movies S4). These parameters, i.e. localization at protruding regions of the cell perimeter and occurrence at essentially immobile membrane sites (Movies S1, S2) set cliffs aside from previously recognized mobile regions of the cells membrane, such as ruffles, lamellipodia, dorsal ridges ^21^. These same structural features were utilized to systematically identify cliff regions in all subsequent investigations.

Cliff membranes were visualized using the signaling super-complex components CD9 and Trop-2 (Figure 3C, Movie S1), whether as endogenous or as transfected molecules. However, in MTE4-14 cells cliffs were also revealed in parental cells devoid of Trop-2, supporting a model of cliffs as broad containers/activation sites of heterogeneous classes of cell growth inducers, that did not depend on Trop-2 expression. Cliff membranes were shown to envelope a rim of cytoplasm, which was easily visualized by calcein and GECO Ca^2+^ probes, CFSE or free cytoplasmic GFP and mCherry, or by cytoplasmic signal transducers (Grb-2-GFP, P-PKCα) (Figure 3). Parallel evidence in MDA-MB-231 and MCF-7 human breast, DU-145 prostate cancer cells and in transformed murine thymus MTE4-14 cells (Figure 1) suggested conservation of cliff features across transformed cell histotypes.

Landscape portraits of membrane cliffs were obtained in living cells through confocal X,Y,Z,T-stacks of CD9–mCherry transfectants. Cliff height and width were found to recursively fluctuate over time at the same sites of the cell membrane (Movies S1, S5). CD9 density was correspondingly found to oscillate over time (Movie S5, pseudocolor representation), first suggesting that recruitment of protein signal transducers at cliffs followed a cyclic behavior.

### Lipid-normalized signal transducer protein content of membrane cliffs

To tackle quantitative structural-functional issues in cliff generation and dynamics we took advantage of previously developed fluorescence quantification and imaging technologies ^15,22–30^. To prevent quantitative artifacts due to the inclusion of different membrane volumes in different confocal sections or cell perimeter sites, we normalized signal transducer density versus membrane lipid content, using diverse classes of fluorescent lipophilic tracers. These included 3,3’-dioctadecyloxacarbocyanine perchlorate (DiOC, **18-carbon chain**); 4-(4-(dihexadecylamino)styryl)-*N*-methylpyridinium iodide (DiA, also known as 4-Di-16-ASP, **16-carbon chain**); 1,1’-didodecyl-3,3,3’,3’-tetramethylindocarbocyanine perchlorate (DiIC, **12-carbon chain**). DiOC, DiA, and DiIC selectively partitioned into different lipid phases of the cell membranes. DiA induced rapid toxicity in living cells and was thus abandoned. Composite parametrization of DiOC and DiIC confocal microscopy signals showed the lowest variance across cell perimeter determinations, and was used to obtain normalized lipid/protein ratios (Table S1).

Lipid-normalized CD9 protein signals were shown to be largely homogeneous throughout the cell perimeter, with the exception of spikes of signaling protein density at cliffs (149 cliff segments; 9-14 segments per individual cliff), indicating preferential recruitment of protein signal transducers at these sites (Table S1). Corresponding findings were obtained from 3D reconstructions and Z-stack analysis of cell populations assessed for endogenous signal transducers and of FP chimera-transfected living cells.

### Cliffs are signaling platforms

We had previously shown that Trop-1 and Trop-2 transduce Ca^2+^ signals upon cross-linking with anti-Trop monoclonal antibodies (mAb) ^14,15^. We thus assessed whether cliffs function as *bona fide* signaling platforms, i.e. as scaffolds for co-localizing signal transducers, thus facilitating their interaction ^19,20^. This was explored in endogenous Trop-2-expressing breast MDA-MB-231 and in thymus MTE4-14/Trop-2 transfectants. After addition of 162-46.2 anti-Trop-2 mAb to transformed cells, Ca^2+^ signaling waves were shown to originate from membrane cliffs (Movie S6, Figure 3B), with a latency time of 7.5±1.0 s (mean±SEM; range: 3.9-15.7 s). The Ca^2+^ signals lasted 81.2±12.1 s (range: 31.4-141.5 s). No signal was elicited in control cells devoid of Trop-2. Trop-2-induced Ca^2+^ signals were shown to lead to activation of PKCα, which functions as a pivotal inducer of Trop-2-induced cell growth ^14^. Recruitment of PKCα-EGFP, a fully functional growth-driving molecule, was found to only occur at cliffs (Figure 2B, Movie S7). PKCα-EGFP membrane recruitment occurred with a latency time of 5.0±0.5 sec (range: 3.9-7.9 sec) after addition of anti-Trop-2 mAb, and lasted on average 55.0±10.6 sec (range: 7.9-125.8 sec). This closely corresponds to overall values for PKCα membrane transport/activation ^14^, consistent with a model whereby cliffs function as main platforms for PKCα signaling.

### Space dimensions of cliffs

The spatial features of cliffs were analyzed on human breast MDA-MB-231 and on murine transformed MTE4-14 cells, over ∼50,000 independent confocal microscopy images (Figure 4). Distribution analysis of these findings showed a reproducible mean cliff length of 27±8 μm across the different cell types analyzed (Table S2). Hence, cliffs appear orders of magnitude larger than previously recognized cell membrane signaling platform in living cells ^31–36^.

**Figure 4.**
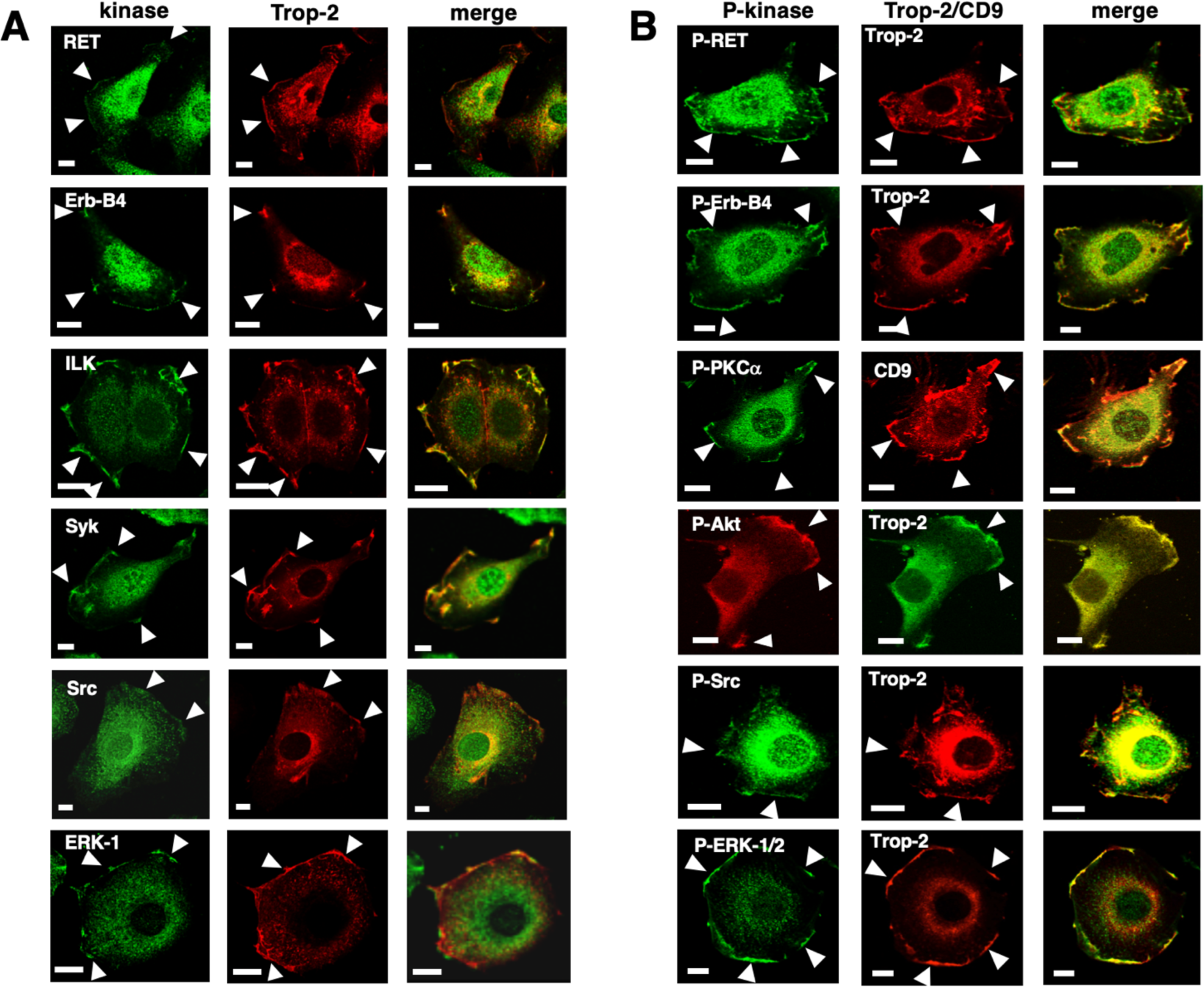
Growth-inducing kinases are recruited and activated at cliffs. Endogenous cytoplasmic kinases were analyzed by mAb-staining immunofluorescence confocal microscopy. Trop-2 and CD9 were utilized as cliff tracers and for colocalization analysis. Representative single-plane images of Z-stack reconstructions are shown. Arrowheads indicate areas of co-recruitment of pairs of signal transducers at membrane cliffs. **(A)** MTE4-14/Trop-2 transfectants were stained for Trop-2 (*red*) and for RET, Erb-B4, ILK, Syk, ERK-1, as indicated (*green*). Scale bars, 10 µm. (**B**) MTE4-14/Trop-2 transfectants were stained for Trop-2 or endogenous CD9 and for the activated/phosphorylated P-Src, P-ERK-1/2, P-Erb-B4, P-RET, P-PKCα, P-Akt as indicated. Scale bars, 10 µm.

### Cliffs are long-lived, recursive platforms

Cliff lifetimes were investigated in living MTE4-14 cells transfected with signaling protein-FP chimeras, capturing images at 30-96 sec intervals (308 cliff acquisitions; 20-37 frames per individual cliff) (Figure 4B). This indicated cliff lifespans of 226±40 sec (mean±SEM) (range: 30-940 sec). Corresponding findings were obtained using shorter image acquisition times (1.18 sec/frame) under continuous-recording mode, which prevented artifacts due to movement-related, out-of-focus shifting of the membrane regions under analysis. Thus, cliff lifetimes appear orders of magnitude longer than those of previously reported signaling platform in living cells ^32,33^.

As indicated above, cliffs recursively assemble at conserved sites of the cell membrane perimeter (Movies S1, S2, Table S2). Wave periods, i.e. the interval between sequential peaks of protein density at cliffs (Figure 4, Table S2), were 293±46 sec (mean±SEM) (12 sequential cycles; range: 70-600 sec). Wave periods thus appeared in the range of cliff lifetimes, suggesting that the recruitment rates of signal-transducers at the cell membrane drive cliff assembly periodicity. These findings were confirmed by high frame-rate/continuous-mode recording confocal microscopy.

Approximately half of the cliffs were found to completely disappear, to then reappear in a cyclic manner, while the remaining cliffs oscillated between maximum and minimum dimensions. The ratio between the maximum and minimum size of individual cliffs in the latter subset appeared centered around 5-fold (median: 4.94; range: 1.99-15.74; near-median measurements: 10 out of 18 cases) (Table S2), suggesting fine regulation of cliff size over time. The minimum fraction of cell membrane perimeter occupied by cliffs at any one time was !5% (Table S2i-ii).

### Cell growth-driving kinases are recruited at cliffs

Trop-1 and Trop-2 stimulate cell growth and tumor progression through downstream cytoplasmic kinases ^2,11,14,16,17^. We thus explored whether cliffs were recruitment sites for Trop-activated kinases. PKCα-FP ^14^ and ERK-FP were shown to dynamically colocalize with Trop-1/Trop-2 at cliffs (Movie S2, Figure 2B). Endogenous growth-driving kinases followed corresponding patterns. We utilized to this end 43 different antibodies from independent suppliers, whenever possible as pairs of independent antibodies for detecting individual target proteins (Material and methods) ^11^. In cases of transfected proteins, e.g. Trops, tetraspanins, care was taken to avoid overexpression of the transfected plasmids, via selection of expressing cells by flow cytometry for the average levels of expression detected in cancer cells ^14^. Following these procedures, we reproducibly detected endogenous Ret, Erb-B4, ILK, Akt ^16^, Src, Syk, ERK-1 at Trop-2/CD9 sites in MTE4-14/Trop-2 cells (Figures 4C, 5A). We went on to show that the recruited kinases were phosphorylated at kinase activation sites under baseline conditions (P-Src, P-ERK-1/2, P-Erb-B4, P-RET, P-Akt, P-PKCα, Figures 4C, 5B), indicating cliffs as platforms for kinase function activation. Consistently, higher levels of kinase activation at cliffs were shown to be induced by treatment with growth factors (GF) (see below).

**Figure 5.**
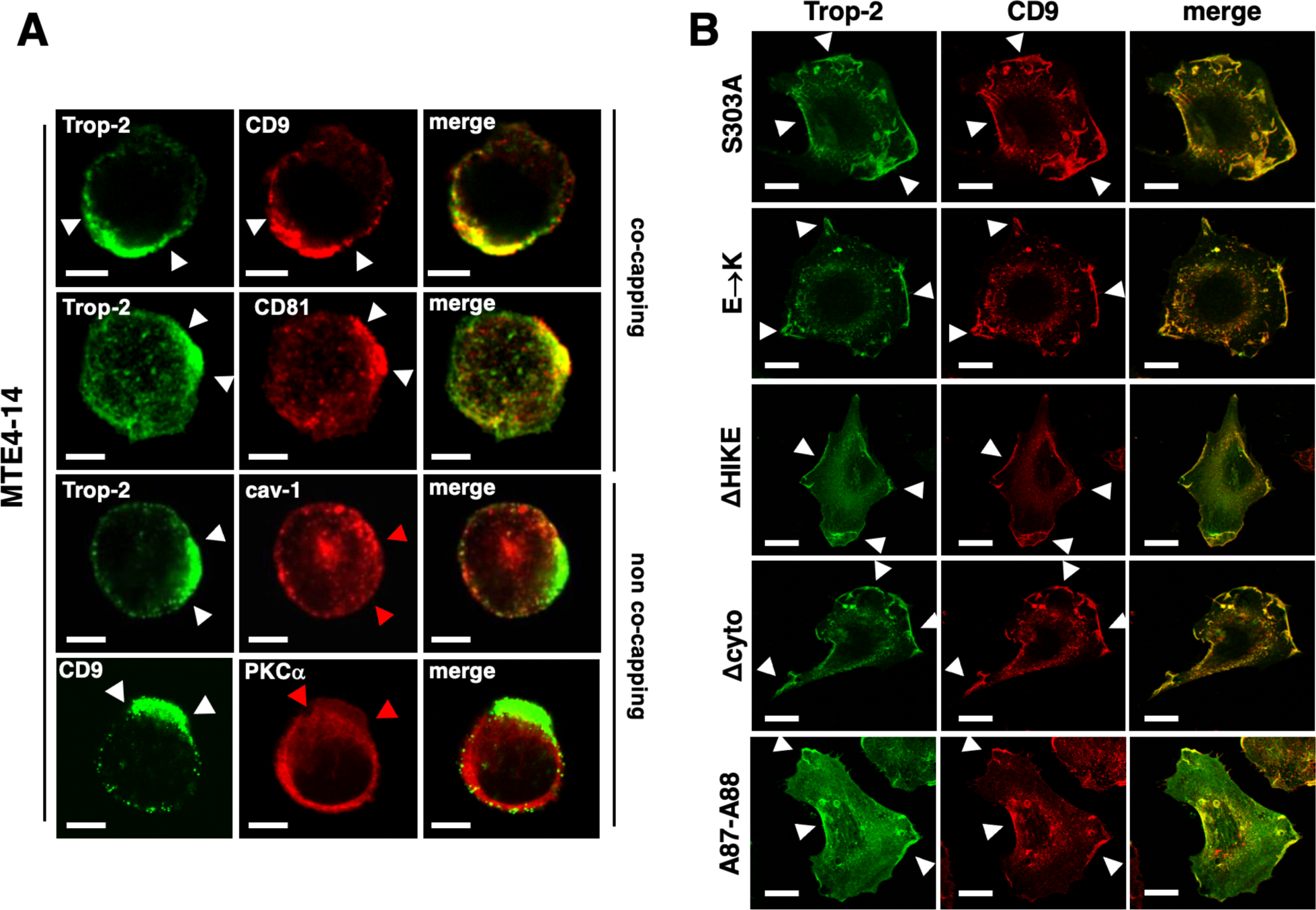
Selective co-recruitment of signal transducers at cliffs. (**A**) (*top*) Co-capping of Trop-2 with CD9 and CD81 in MTE4-14 cells. (*bottom*) Lack of Trop-2 co-capping with caveolin-1 and control PKCα. White arrowheads indicate the edges of the capped regions. Red arrowheads indicate the regions with lack of co-capping. Bars, 5 µm. (**B**) White arrowheads indicate colocalization of Trop-2 and CD9 at cliffs. Trop-2 mutants were generated as described ^14^. Mutants of the cytoplasmic region included: S303A: mutation of the PKCα phosphosite. E->K: mutation of the four E in the cytoplasmic tail to K. ΔHIKE: deletion of the HIKE region. Δcyto: deletion of the cytoplasmic tail. A87-A88: R87A and T88A mutants of the ADAM10 extracytoplasmic cleavage site ^10^. None of the tested mutants detectably affected Trop-2 recruitment at cliffs. Bars, 5 µm.

Comparative analysis of absolute levels of expression and phosphorylation status of receptor tyrosine kinases (RTK) and of cytoplasmic kinases at cliffs, as compared to non-cliff sites, indicated that the vast majority of activated/phosphorylated kinases did associate to membrane cliffs (99% of total membrane content for P-Erb-B4; 91% for P-Ret; 90% for P-PKCα; 90% for P-Src) (Table S3), suggesting cliffs as main kinase activation sites at the plasma membrane.

PKCα is recruited to the cell membrane upon Ca^2+^ signaling (Movies S6, S7), and is subsequently activated by phosphorylation at S657 ^14^. We thus asked whether recruitment at cliffs preceded PKCα activation. We tackled this issue by assessing whether PKCα kinase activity was required for cliff localization or was dispensable. We found that cliff recruitment was kinase activity– independent, as a dominant-negative, kinase-inactive K368R PKCα-GFP ^37^ was efficiently recruited at cliffs (Movie S8).

We asked a corresponding question for Trop-2 recruitment at cliffs. Multiple inactive Trop-2 mutants were generated ^14^. These included S303A, a mutation of the acceptor site of phosphorylation by PKCα, which prevents PKCα recruitment at cliffs. Deletion of the HIKE region (ΔHIKE), a regulatory site for protein-protein and protein-phospholipid interactions ^38,39^, correspondingly prevented Trop-2 from recruiting PKCα to the cell membrane, despite preserving Trop-2 induction of intracellular Ca^2+^ waves ^14^. Trop-2 mutants of four cytoplasmic tail E to K (E->K) and deletion-mutants of the entire cytoplasmic tail (Δcyto) were correspondingly assessed. None of the tested mutants detectably affected Trop-2 recruitment to cliffs (Figure 6), first suggesting that functional signaling of Trop-2 was not required for recruitment at cliffs.

**Figure 6.**
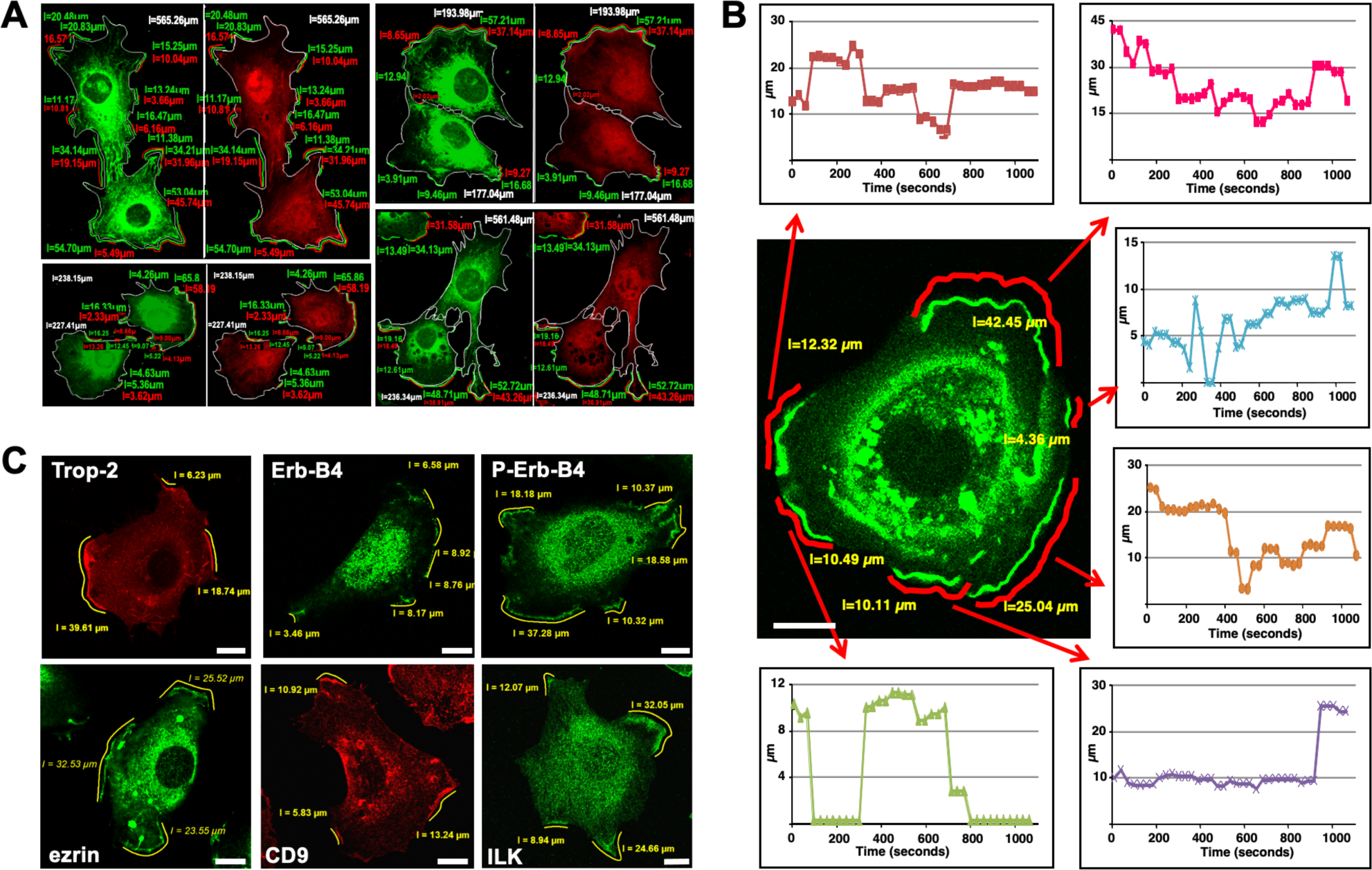
Space and time dimensions of membrane cliffs. (**A**) Analysis of membrane cliff size. Yellow stripes: overlaid ‘rulers’ were generated by the LSM Image Browser 4.0 software. Cliff length (µm) was estimated by combining the overlay and measure routines. (**B**) Time-lapse analysis of membrane cliff size over time. The full data-set is presented in Table S2. Analysis of cliff space-time transitions was performed on individual MTE4-14 living cells transfected with a Trop-2-EGFP chimera. Movies were recorded using the Zeiss LSM 510 3.0 software. Pixel residence time was set at 1.60 µs to 3.10 µs; image format was 1024×1024 pixels at 8 bit pixel depth. Images were captured at 30 s to 96 s intervals. Individual platform lengths (red stripes) are plotted as graphs versus time. Bar, 5 µm. (**C**) Cliff size was estimated for membrane-recruited Trop-2, CD9, Erb-B4, phospho-Erb-B4 (P-Erb-B4), ezrin, ILK. Measured cliff length in µm is indicated. Bars, 5 µm.

Cytoplasmic tail mutagenesis (S303A, ΔHIKE, Δcyto) abolishes the capacity of Trop-2 to induce cell growth ^10,14^. Extra-cytoplasmic cleavage of Trop-2 by ADAM10 is required to activate Trop-2 as a growth inducer ^10^. We thus explored whether Trop-2 recruitment at cliffs was affected by mutagenesis of the ADAM10 cleavage site (R87A-T88A ^10^). R87A-T88A Trop-2 mutants were efficiently transported at cliff sites (Figure 6), albeit unable to signal for cell growth ^10^, supporting the notion that recruitment at cliffs does not require Trop-2 activation.

### Cliff signaling underlies induction of cell growth

Trop-2 drives signaling for cell growth and shRNAs for *TROP2* abolish the growth of Trop-2-expressing cells ^14^. Correspondingly, shRNA inhibition of endogenous Akt ^16^, CD9 or PKCα ^14^ prevent Trop-2 signaling for growth, revealing a tight interplay of signaling super-complex components that are co-recruited at cliffs. The actin cytoskeleton informs EGF uptake and modulates EGF receptor activation and downstream signaling ^40^. Activated RTK and Ser/Thr protein kinases are engaged at cliffs, leading us to challenge a broader model, i.e. whether GF induce cliff assembly, to then induce cell growth. Fetal calf serum/growth-factor deprivation of parental MTE4-14 cells was shown to lead to the progressive disappearance of cliffs, and to a parallel reduction of endogenous P-PKCα and CD9 membrane levels after 24 h to 48 h starvation, and to the arrest of cell proliferation (Table S3, Figures 7, S2).

**Figure 7.**
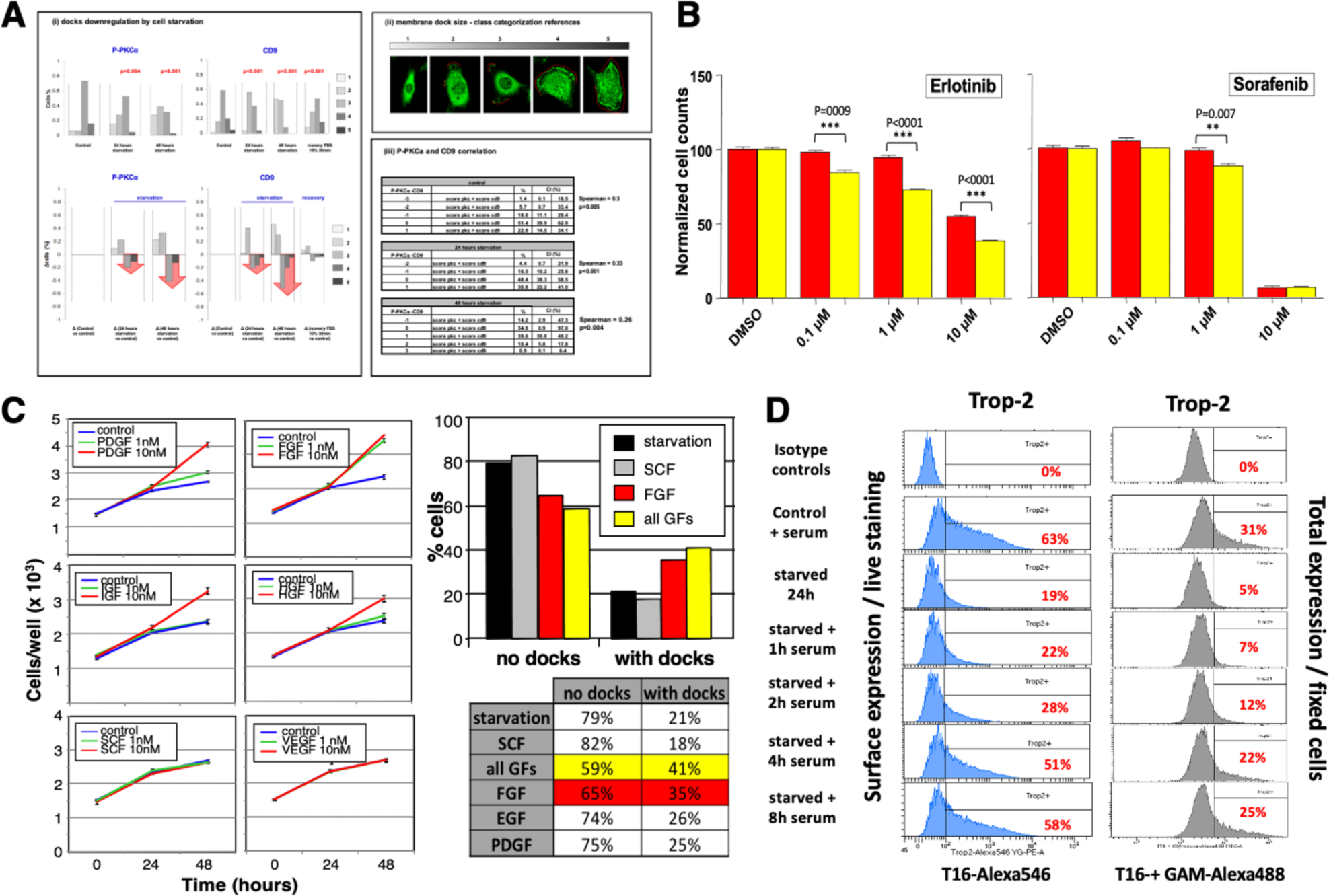
Membrane cliff and cell growth induction by GF. (**A**) MTE4-14 cells were growth factor-starved (0.1% serum) for 24 h or 48 h. After treatment, cells were fixed and stained. Co-recruitment of endogenous P-PKCα and CD9 at cliff sites was quantified by image analysis. (**i**) (*top*) Cells were classified (classes 1-5) according to the levels of expression of p-PKCα or CD9 at cliffs. (*bottom*) Histograms of differential class distributions of treated versus control cells. (**ii**) Intensity scores were computed by multiplying perimeter values of regions containing P-PKCα (length of cliffs; highlighted in red) (5=entire cell perimeter; 1=no platforms) to intensity values (5=highest intensity; 1=undetectable); the square root of the product was used for cell categorization. (*top right*) representative cells for the five perimeter score classes are shown. (**i**, *bottom panel*; **iii**) Correlation between p-PKCα and CD9 across the different groups. To highlight score trends, the score of CD9 was substracted from the score of p-PKCα. Results were then classified as above or below zero. Spearman’s rank correlation coefficients and P values are indicated. The full datasets and comparison statistics are presented in Table S3. **(A)** P-PKCα and CD9 membrane localization levels upon cell starvation and serum-induced recovery. (**B**) Inhibition of cliff-induced cell growth by kinase inhibitors. Scale bars: normalized cell/well numbers at 72 hours after seeding. Red: Trop-2 transfectants. Yellow: vector-alone transfectants. P values (ANOVA) of Trop-2 versus control cells are indicated; (*left*) Erlotinib as EGFR ^41^, FGFRs ^42^ inhibitor; (*right*) Sorafenib as VEGFRs inhibitor ^41,42^. Trop-2 expression antagonized cell growth inhibition by Erlotinib, but not that by Sorafenib (dose-response: 0.1 µM to 10 µM). (**C**) (*left*) Growth curves of serum-starved MTE4-14 cells, rescued by treatment with GF. PDGF and FGF-1 efficiently induced cliffs and cell growth. HGF, IGF-1 showed lesser impact on both cliffs and cell growth, SCF, VEGF had no significant effect on either parameter. (*right*) Percentage of cells with or without membrane cliff formation, as induced by 24 h exposure to the GF indicated, 10 nM. Cliffs were visualized by staining for P-PKCα. The largest induction was by the mixture of EGF, FGF, PDGF (‘all GFs’) (in 41% of cells). The largest induction by a single GF was by FGF (in 35% of cells). (**D**) GF-induced recovery of Trop-2 synthesis and transport to the cell surface. Cells in culture were starved in 0.1% serum for 24 h. Serum addition triggered both Trop-2 synthesis (gray profiles; total Trop-2 in fixed cells) and Trop-2 transport to the cell surface (blue profiles; cell membrane-only staining of live cells). The slope of global synthesis of Trop-2 and transport to the cell membrane over time indicated faster transport recovery versus synthesis. Progressive recovery of membrane Trop-2 levels was reached after ≥8 h serum treatment.

Addition of serum to serum-starved HCT116 colon cancer cells led to the recovery of Trop-2 synthesis and transport to the cell surface after 1 h exposure of cells starved for 48 h. Trop-2 levels subsequently reached those in control cells in ≈8 hours (Figure 7D). Purified FGF-1, PDGF, EGF, HGF, IGF-1, SCF and VEGF were next tested for their capacity of inducing membrane cliffs in serum-starved cells and for driving cell growth. A 30 min stimulation with 1, 10 or 100 nM PDGF, FGF-1 or EGF sufficed to induce cliff formation and cell growth (Figures 7C, S2). HGF, IGF-1 showed a lesser impact, whereas SCF, VEGF had essentially no effect, on both cliff formation (Table S3B,C) and on cell growth (Figures 7C, S2), supporting a model whereby cliff formation by GF/disruption by GF deprivation parallel induction/ablation of cell growth, respectively. Trop-2 expression was shown to interact with CD9 expression levels and with response to GF (Figure S2B). Consistent with the GF specificity of cliff induction and of cell growth triggering, Trop-2 expression was shown to antagonize, in a dose-response manner, cell growth inhibition by 0.1 µM to 10 µM Erlotinib, which largely acts as EGFR ^41^ and FGFR1-4 ^42^ inhibitor. On the other hand, Trop-2 did not antagonize cell growth inhibition by Sorafenib, which largely acts as Ser/Thr CDK and Tyr VEGFR1-3 inhibitor ^41,42^ (Figure 7B).

Thus, specific GF induce formation of membrane cliffs. A mixture of FGF-1, PDGF and EGF had a stronger impact than any GF alone, and approximately doubled both the number of cliffs observed in cells grown in conventional serum-supplemented medium and cell proliferation (Figure 7C), suggesting that multiple growth factors exert an additive effect on cliff assembly and signaling. An extended analysis is presented in Table S3B, whereby CD9/P-PKCα cliff signal intensity versus occupied perimeter fraction versus growth factor stimulation was performed on a cell-by-cell basis, on ≈2440 cells, across 122 image panels, recorded as correlated pairs of fluorescence channels, and on additional ≈1680 cells, across 84 single-fluorescence validation panels.

These findings highlighted distinct time-scales for the periodic assembly/disassembly of cliffs during steady-state conditions (∼200 sec), versus *de novo* formation as induced by GF (1-8 hours). The transport of Trop-2 to the cell surface was measured after starvation/refeeding (flow cytometry analysis of living HCT-116 cells). Cell surface Trop-2 was detectable after one hour of serum re-feeding in 22% of the cells, but reached baseline levels in the majority of the cells in !8 hours (Figure 7D). Trop-2 neosynthesis, as measured by subtracting membrane-only staining from whole cell signals, was shown to account for a considerable fraction of recovery kinetics, indicating that there is a requirement for signal transducer neosynthesis during cliff recovery after starvation, which significantly extends the kinetics of cliff generation, versus baseline control conditions.

### The β-actin cytoskeleton informs cliff dynamics

Disruption of actin microfilaments, but not of microtubuli or intermediate filaments, was shown to affect the localization of Trop-1/Ep-CAM at cell-cell boundaries ^43^. Our findings showed that activation of the Trop-2 membrane super-complex remodels the β-actin/α-actinin cytoskeleton through cofilin-1, annexins A1/A6/A11 and gelsolin ^14^. We show here that co-recruitment of ezrin, moesin, fascin, α-actinin, cortactin, vinculin and Trop-2–GFP occurs at cliffs (Figure 8, S3). Co-transfection of MTE4-14 cells with the β-actin cytoskeleton marker Lifeact–mRFP1 and Trop-2– GFP showed extensive co-recruitment of actin cytoskeleton and Trop-2–GFP at cliffs (Movie S9). Analysis of endogenous molecules through phalloidin-FITC for β-actin and mAb-Alexa633 for Trop-2 showed a corresponding colocalization at cliffs in fixed MTE4-14 cell transfectants (Figure S3). Crosslinking of Trop-2 in living cells using the 162-46.2 mAb induced β-actin polymerization in parallel to Ca^2+^ (Movie S10) and PKCα signaling (Movie S7). Induced β-actin polymerization showed a latency time of 10.2 ±2.9 sec (mean±SEM; range: 3.9-19.7 sec) and a plateau after 55.0 ±13.5 sec (range: 23.6-90.4 sec).

**Figure 8.**
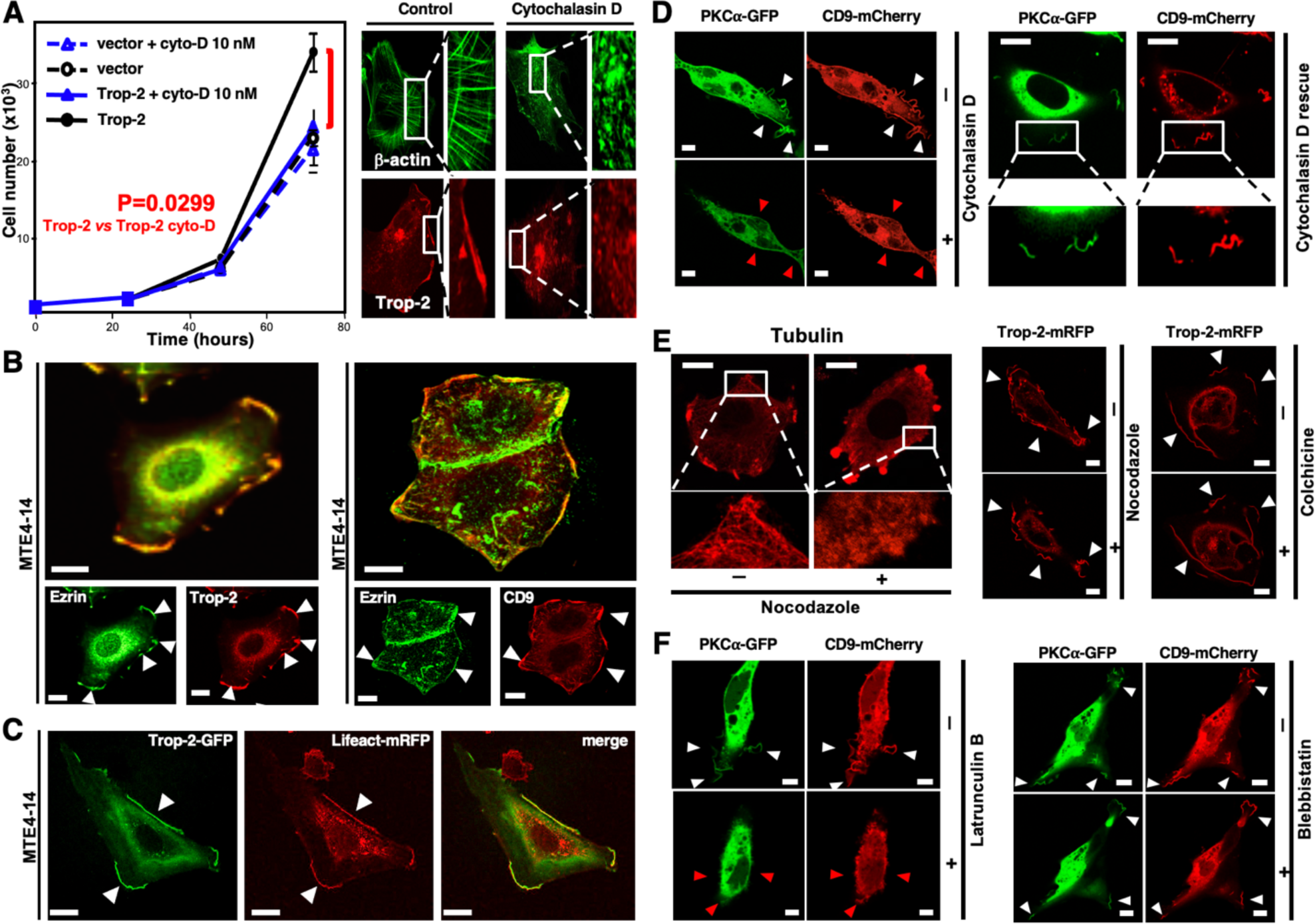
β-actin informs cliff structure and function. MTE4-14 cells were analyzed for colocalization of signal transducers with cytoskeleton components in resting cells and upon disruption of β-actin (cytochalasin D, latrunculin B), tubulin (colchicine, nocodazole), myosin (blebtatin) assemblies. Signal transducer colocalization at cliffs is indicated by white arrowheads. Red arrowheads indicate no colocalization of signal transducers or loss of it upon cytoskeleton disruption. (**A**) Growth curves of MTE4-14 cells transfected with Trop-2 or empty vector, treated with 10 nM cytochalasin D (cyto-D). Growth curves were compared by two-way ANOVA, with Bonferroni correction. Data are presented as mean±SEM. (*right*) Phalloidin-FITC-staining of β-actin (*top*, green) and Alexa633-mAb staining of Trop-2 (*bottom*, red) show loss of β-actin polymerization and of membrane cliffs upon treatment with cytochalasin D. (**B**) Trop-2 and CD9 colocalize with ezrin at cliffs. Scale bars, 10 µm. (**C**) Co-transfection of MTE4-14 cells with the β-actin cytoskeleton marker Lifeact–mRFP1 and Trop-2–GFP showed extensive co-recruitment of actin cytoskeleton and Trop-2–GFP at cliffs. The full dataset is presented in Movie S4. Scale bars, 20 µm. (**D-F**) Colocalization of PKCα–GFP and CD9–mCherry in MTE4-14 cell transfectants was explored in resting cells and upon cytoskeleton disruption. (**D**) (*left*) Cytochalasin D treatment. (*right*) Recovery of membrane platforms 24 h after removal of cytochalasin D. (**E**) Disruption of tubulin organization after treatment with nocodazole (*left*) or colchicine (*right*) had no effect on cliffs. Tubulin was stained with anti-tubulin mAb-Alexa546. (**F**) (*left*) Latrunculin B treatment disrupted PKCα–GFP/CD9–mCherry colocalization at cliffs. (*right*) Blebbistatin treatment had no impact on cliff structure. Scale bars, 5 µm.

We thus assessed whether drug-induced depolymerization of the β-actin cytoskeleton affected cliff formation. At variance with tetraspanin microdomains ^44,45^, treatment with latrunculin B or cytochalasin D (which bind to G-actin and prevent polymerization) led to the disappearance of PKCα-GFP and CD9-mCherry-hosting membrane cliffs (Figure 8, Movie S11). Dose-response treatments of MTE4-14/Trop-2 transfectants revealed that 10 nM cytochalasin D sufficed to disrupt cliffs and to revert the growth rate of Trop-2 transfectants to the growth rate of control cells (Figure 8A). Cliffs and Trop-2-induced cell growth were rescued after washout of cytochalasin D, in parallel to the recovery of the actin cytoskeleton (Figure 8D).

Vesicular traffic of growth-regulatory receptors to the cell surface depends on microtubules, which are orchestrated by merlin through a Rac/MLK/p38 (SAPK) pathway ^46^. Tubulin-binding inhibitors of microtubule polymerization (nocodazole, colchicine) efficiently inhibited microtubule assembly. However, they had no detectable impact on cliff formation (Movie S12, Figure 8E), overall suggesting no involvement of the microtubule cytoskeleton in cliff scaffolding.

Membrane recruitment of myosin IIa–mTFP1 was detected at later times points (38.9 ±7.6 sec; range: 11.8–70.7 sec) than that of polymerized β-actin, and lasted for considerably longer times 179.5 ±7.7 sec (range: 161.1-192.6 sec). Consistent with a distinct functional role of myosin versus β-actin in the generation of cliffs, the myosin inhibitor blebbistatin had no influence on cliff formation (Figure 8F). Taken together, these findings indicated that cliff formation distinctly depend on the β-actin cytoskeleton, and that cliff disappearance and loss of cliff signaling capacity is not an obligate, nonspecific response to any cytoskeletal perturbation.

## Conclusions

Our findings indicate cliffs as novel, macroscopic signaling platforms, that recruit and activate drivers of cell growth. Cliffs were induced by GF, acted as sites of kinase recruitment/phosphorylation/activation and as sites of origin of Ca^2+^ signaling. Signaling competence for cell growth was acquired through the co-recruitment of trans-membrane (RTK, CD9, CD81, CD98, Co-029, CD316, Trop-1, Trop-2), and cytoplasmic (PKCα, ERK, Akt, Src, Syk, ILK, ezrin) signal transducers to cliffs. On the other hand, disruption of cliff assembly by GF deprivation or by β-actin depolymerization abolished signaling for cell growth. Loss-of-function mutagenesis of Trop-2 and PKCα did not prevent their recruitment at cliffs, suggesting a model whereby inactive signaling molecules are recruited at cliffs, for subsequent activation by specific interactors.

Cliffs were shown to be ≈27 μm-long segments of the cell membrane, that recursively formed over hundreds of seconds at conserved sites of the cell perimeter. Thus, cliffs appear orders of magnitude larger than any previously recognized cell membrane signaling platform in living cells ^31–36^. Cliff lifetimes correspondingly appeared orders of magnitude longer than those of any previously reported signaling platform in living cells ^32,33^, suggesting cliffs as high-dimensional signaling platforms for cell growth in normal, living cells ^31,47,48^.

### Box 1

- Cliff are macroscopic, recursive cell membrane platform that trigger signals for cell growth.
- Cliffs are two-leaflet membrane elevations, up to 1.5 µm tall, ≈27 µm wide.
- Cliffs are long-lived membrane platforms 226±40 sec (mean±SEM) (range: 30-940 sec).
- Cliffs recur over long time frames 293±46 sec (mean±SEM) (12 sequential cycles; range: 70-600 sec), at essentially immobile sites of cell edges.
- Cliffs are co-recruitment activation sites of multiple classes of signal transducers for cell growth.
- Cliffs are sites of phosphorylation/activation of growth-inducing protein kinases and are trigger sites of calcium waves.
- Cliffs are induced by growth factors and disappear with serum starvation.
- Cliffs selectively depend on the β-actin cytoskeleton and β-actin depolymerization abolishes cliff signaling for cell growth.

## Acknowledgments

We thank I. Upmann and M. Garcia-Parajo for support with NSOM microscopy. We thank J. Schlessinger, K. Simons, H. Morrison, R. Plebani, L. Antolini, K. Havas, D. Parazzoli, A. Oldani, A. D’Angelo and G. V. Beznoussenko for help and discussion during this work. We gratefully acknowledge the support of Fondazione Cassa di Risparmio della Provincia di Chieti, Fondazione Compagnia di San Paolo (Grant 2489IT), the Ministry of the University and Research (MIUR, Italy) (SCN_00558), the Ministry of Development (MISE, Italy) (MI01_00424), Region Abruzzo (POR FESR 2007-2013: Activity 1.1.1 line B), the Italian Association for Cancer Research (AIRC, Italy), the Programma Per Giovani Ricercatori “Rita Levi Montalcini” (MIUR, Italy – Grant PGR12I7N1Z for support to M.T, the European Regional Development Fund (FEDER) “Una manera de hacer Europa” for support to V.R.C. and M.Z.. FLIM and STED were performed at the Centro National de Investigaciones Cardiovasculares, CNIC (Madrid, Spain), which is supported by the Instituto de Salud Carlos III (ISCIII), the Ministerio de Ciencia e Innovación (MCIN) and the Pro CNIC Foundation.

The sponsors had no role in the design and conduct of this study, nor in the collection, analysis, and interpretation of the data, nor in the preparation, review, or approval of the manuscript.

