## Supplementary online for "Membrane cliffs are giant, recursive platforms that drive calcium and protein kinase signaling for cell growth"

**Supplementary Information**

Table of contents

FIGURES S1-S3

Figure S1. NSOM analysis of Trop-1 and Trop-2 at cliffs.

Figure S2. Growth factors induce membrane cliffs.

Figure S3. The β-actin cytoskeleton scaffolds membrane cliffs.

TABLES S1-S3

Table S1. Lipid versus signal transducer protein content at membrane cliffs.

Table S2. Space and time parameters of cliff recursive assembly.

Table S3. Growth factors-driven recruitment and activation of signaling kinases at membrane cliffs.

MOVIES S1-S12

Movie S1. Co-recruitment of CD9 and Trop-2 at cliffs.

Movie S2. Recursive co-recruitment of Trop-2 and ERK at cliffs.

Movie S3. 3D reconstruction of cell membrane cliffs.

Movie S4. Cliff occurrence is unrelated to cell movement.

Movie S5. Four-dimension cliff dynamics.

Movie S6. Signal triggering induces $Ca^{2+}$ waves from cliffs.

Movie S7. PKCα signaling is induced at cliffs.

Movie S8. Kinase-inactive PKCα recruitment at cliffs.

Movie S9. Co-recruitment of the β-actin cytoskeleton and Trop-2 at cliffs.

Movie S10. Assembly of the β-actin cytoskeleton at cliffs parallels induction of $Ca^{2+}$ signaling.

Movie S11. Integrity of the β-actin cytoskeleton is required for membrane cliff scaffolding.

Movie S12. Microtubule integrity is not required for membrane cliff scaffolding.

EXPERIMENTAL PROCEDURES
Cell lines and reagents
Cell biology and biochemistry
Microscopy
Immuno-electron microscopy
STATISTICS

REFERENCES

### STAR ★ METHODS

### KEY RESOURCE TABLE

| REAGENT or RESOURCE | SOURCE | IDENTIFIER |
|---|---|---|
| **Antibodies** | | |
| Trop-2 | ATCC | 162-46.2/HB-187 |
| Trop-2 | [1] | T16 |
| β-tubulin | Sigma-Aldrich | TUB 2.1 |
| CD9 | Santa Cruz Biotech | sc-18869 |
| CD9 | Abnova | 14-0098 |
| CD44 | Santa Cruz Biotech | sc-52536 |
| CD98 | Abnova | 14-0829-80 |
| CD151 | Santa Cruz Biotech | sc-18753 |
| CD151 | Abnova | H00000977-M02 |
| Co-029 | Gift from F. Lanza | AZM 22.2 |
| ERK1 | Santa Cruz Biotech | sc-94-G |
| EGFR | Santa Cruz Biotech | sc-03-G |
| caveolin-1 | Santa Cruz Biotech | sc-894 |
| phospho-ERK1/2 (pT177/pT160) | Santa Cruz Biotech | sc-23759-R |
| Ret | Santa Cruz Biotech | sc-13104 |
| phospho-Ret (pY1062) | Santa Cruz Biotech | sc-20252-R |
| ErbB-4 | Santa Cruz Biotech | sc-283 |
| Syk | Santa Cruz Biotech | sc-1077 |
| c-Src | Santa Cruz Biotech | sc-19 |
| phospho-c-Src (pY144) | Santa Cruz Biotech | sc-12928-R |
| ezrin | Santa Cruz Biotech | sc-20773 |
| phospho-ezrin (pY354) | Santa Cruz Biotech | sc-101678 |
| NF2/Merlin | Santa Cruz Biotech | sc-332 |
| phospho-Akt (pS473) | Cell Signaling Tech | 736E11 |
| phospho-PKCα (pS657) | Santa Cruz Biotech | sc-12356 |
| PKCα | Santa Cruz Biotech | sc-208 |
| CD81 | GeneText | GTX75430 |
| phospho-ErbB-4 (pY1162) | Epitomics | 2295-1 |
| ILK | Millipore | EPR1592 |
| EphB2 | Santa Cruz Biotech | sc-130068 |
| AlexaFluor 488 goat anti-rat IgG | Life Technologies/Molecular Probes | A-11006 |
| AlexaFluor 633 goat anti-rat IgG | Life Technologies/Molecular Probes | A-21094 |
| Alexa Fluor 488 goat anti-rabbit IgG | Life Technologies/Molecular Probes | A-11008 |
| Alexa Fluor 633 goat anti-rabbit IgG | Life Technologies/Molecular Probes | A-21070 |
| Alexa Fluor 488 donkey anti-goat IgG | Life Technologies/Molecular Probes | A-11055 |
| Alexa Fluor 633 donkey anti-goat IgG | Life Technologies/Molecular Probes | A-21082 |
| Alexa Fluor 488 goat anti-mouse IgG | Life Technologies/Molecular Probes | A-11001 |
| Alexa Fluor 633 goat anti-mouse IgG | Life Technologies/Molecular Probes | A-21052 |
| HRP anti mouse | Calbiochem | 401215 |
| HRP anti rabbit | Santa Cruz Biotechnology | sc-2004 |
| HRP anti goat | Santa Cruz Biotechnology | sc-2020 |
| anti mouse for immunohistochemistry | Dako | K4001 (EnVision kit) |
| anti goat for immunohistochemistry | Dako | K0679 (LSAB kit) |
| **Chemicals, Peptides, and Recombinant Proteins** | | |
| Protease inhibitor tablets | Thermo Scientific/Pierce | 88266 |
| Phosphatase inhibitor tablets | Thermo Scientific/Pierce | 88667 |
| Protein G-Sepharose Fast Flow | GE Healthcare | 17-0618-05 |
| DAPI | Thermofisher | D3571 |
| Lipofectamine 2000 | Thermofisher | 11668030 |
| Lipofectamine LTX | Thermofisher | 15338030 |

| | | |
|---|---|---|
| methyl-β-cyclodextrin | Sigma-Aldrich | C4555 |
| 3,3'-dioctadecyloxacarbocyanine perchlorate (DiOC) | Sigma-Aldrich | D275 |
| 1,1'-didodecyl-3,3,3',3'-tetramethylindocarbocyanine perchlorate (DiLC) | Sigma-Aldrich | D383 |
| 4-(4-(dihexadecylamino)styryl)-N-methylpyridinium iodide (DiA; 4-Di-16-ASP) | Sigma-Aldrich | D3883 |
| cytochalasin D | Sigma-Aldrich | C2618 |
| colchicine | Sigma-Aldrich | C9754 |
| blebbistatin | Merck | 203391 |
| latrunculin | Merck | 428020 |
| nocodazole | Merck | 487928 |
| Phorbol 12-myristate 13-acetate ester (PMA) | Sigma-Aldrich | P8139 |
| Calcein AM | Life Technologies | C1430 |
| phalloidin-FITC | Sigma-Aldrich | P5282 |
| phalloidin-TRITC | Sigma-Aldrich | P1951 |
| Mouse EGF | Miltenyi Biotech | 130-094-036 |
| Human FGF-2 | Miltenyi Biotech | 130-093-837 |
| Human HGF | Miltenyi Biotech | 130-093-871 |
| Human IGF-1 | Miltenyi Biotech | 130-093-885 |
| Human PDFG-BB | Miltenyi Biotech | 130-093-980 |
| Mouse SCF | Miltenyi Biotech | 130-094-079 |
| Mouse VEGF | Miltenyi Biotech | 130-094-086 |
| Human EGF | PeproTech | AF-100-15 |
| Erlotinib | SelleckChem | S1023 |
| Sorafenib (BAY 43-9006) | SelleckChem | S7397 |
| **Critical Commercial Assays** | | |
| SuperSignal West Femto ECL Kit | Thermo Scientific/Pierce | 34094 |
| AlexaFluor 488 carboxylic acid, succinimidyl ester | Life Technologies/Molecular Probes | A20000 |
| AlexaFluor 546 carboxylic acid, succinimidyl ester | Life Technologies/Molecular Probes | A20002 |
| AlexaFluor 633 carboxylic acid, succinimidyl ester | Life Technologies/Molecular Probes | A20005 |
| **Experimental Models: Cell Lines** | | |
| MDA-MB-231 | ATCC | N/A |
| MCF-7 | ATCC | N/A |
| HT-29 | ATCC | N/A |
| HCT116 | Roche (Penzberg – Germany) | N/A |
| COLO-205 | ATCC | N/A |
| DU-145 | ATCC | N/A |
| MTE4-14 | P. Naquet | [2] |
| NS-0 | ATCC | N/A |
| **Recombinant DNA** | | |
| pEYFP-N1 | Clontech | 6006-1 |
| pmRFP1 | Gift from R. Tsien | [3] |
| pCFP | Gift from R. Tsien | [3] |
| pmCherry | Gift from R. Tsien | [3] |
| pCMV-SPORT6 | Imagenes | |
| pGFP3-PKCα | Gift from J.W. Soh | [4] |
| pGFP3-PKCδ | Gift from J.W. Soh | [4] |
| **Software and Algorithms** | | |
| Fiji (ImageJ) | | https://fiji.sc/ |
| ZEN 2009 Light Edition | Zeiss | www.zeiss.com |
| VistaVision | ISS Inc., Champaign, IL, USA | www.iss.com |
| SimFCS 4 | Lab. of Fluorescence Dynamics | https://www.lfd.uci.edu/ |
| Sigma Stat 4.0 | Systat Software, Inc. | https://systatsoftware.com |
| SISA | SISA | www.quantitativeskills.com/sisa |
| GraphPad Prism7 | GraphPad Software, Inc. | www.graphpad.com |
| **Other** | | |
| FACS Canto II Cell Analyzer | BD Biosciences | N/A |
| FACS AriaIII Cell Sorter | BD Biosciences | N/A |

| | | |
|---|---|---|
| LSM-510 META confocal microscope | Zeiss | N/A |
| LSM-800 AiryScan confocal microscope | Zeiss | N/A |
| Eclipse-Ti | Nikon | N/A |
| Alba-fast FLIM Workstation | ISS Inc., Champaign, IL, USA | N/A |

### Contact for Reagent and Resource Sharing

#### EXPERIMENTAL MODEL AND SUBJECT DETAILS

**Cell lines**

Human MDA-MB-231 and MCF-7 breast, DU-145 prostate cancer cells were grown in RPMI 1640 medium. Transformed murine thymic epithelial MTE4-14 cells were maintained in Dulbecco's modified Eagle's medium. Complete cell culture media were prepared by adding 10% fetal calf serum (FCS), 100 IU/mL penicillin and 100 µg/mL streptomycin.

**Plasmids**

The enhanced yellow fluorescent protein (EYFP) expression vector was used to generate EYFP chimeric proteins as described previously [5,6]. A corresponding vector devoid of the coding sequence of EYFP (pΔYFP) was used to express Trop-2 in mammalian cells. Wild-type *TROP2* was obtained by PCR from the original full-length *TROP2* clone [7], using the following primers:

Forward (XhoI): 5'-GCGATTctcgagTCCGGTCCGCGTTCC–3'

Reverse (KpnI): 5'-GCGCCggtaccAAGCTCGGTTCCTTTC–3'

The amplified segment was inserted in the vector at the XhoI/KpnI sites.

Chimeric proteins between Trop-2 (XhoI/KpnI), CD9 (EcoRI/BamHI), ERK, EGFR, α-actinin, CD316 (EcoRI/AgeI), and red fluorescent protein (mRFP1), cyan fluorescent protein (CFP), mCherry enhanced green fluorescent protein/ yellow fluorescent protein (EGFP/YFP) were constructed in pΔYFP as above. Wild-type CD9 and CD316 were expressed in the pCMV-SPORT6.

**Antibodies**

Monoclonal antibodies were purified and conjugated with Alexa488, 546 633 as previously described [8]. Rabbit anti-Trop-2 pAb were generated by subcutaneous immunization with recombinant Trop-2 synthesized in bacteria [7]. Trop-2-reactive antibodies were purified by antigen affinity, via binding to recombinant Trop-2 immobilized on NHS-Sepharose columns, and eluted with 0.1 M glycine, pH 2.5.

### METHOD DETAILS

#### Cell biology and biochemistry

*DNA transfection*

Cells were transfected with DNA in Lipofectamine 2000 or LTX (Invitrogen) following manufacturer instructions. Stable transfectants were selected in G418-containing medium.

*Cell-growth assays*

Cells were seeded at 1.5-3.0 $\times 10^3$ cells/well in 96-well plates (five replica wells per data point). Cell numbers were quantified by staining with crystal violet, and normalizing against standard reference curves of two-fold serially diluted cell samples. Crystal violet assays (n=3 per experimental condition) were validated for linearity of response across assay conditions/cell lines versus direct cell counting by image analysis as previously described [9].

MTE4-14 cells transfected with empty vector (control) or Trop-2 were seeded at a density of $10^3$ cells/well in normal medium (untreated) or medium containing cytochalasin D at different concentrations, as indicated [10]. Actual numbers of cells/well at time 0 were measured and used to normalize cell counts for each experimental group.

For cell growth assays after serum-starvation, cells were seeded in complete medium. After 24 h, wells were washed and refed with medium supplemented with 0.1% FCS at time zero. Growth factors (i.e., EGF, FGF-1, HGF, IGF-1, PDGF, SCF, VEGF) were added at 1, 10, 100 nM at 24 h and again at 48 h, as indicated. Cell morphology and cliffs were assessed as indicated below. Erlotinib is an EGFR [11] and FGFR1-4 [12] Tyr kinase inhibitor. Sorafenib acts as a Ser/Thr CDK family inhibitor [11], and as PDGFR $\alpha/\beta$ [11] and VEGFR1-3 family Tyr kinase inhibitor [11,12]. Both inhibitors were dissolved in DMSO and administered at 0.3 $\mu$M, 1 $\mu$M and 3 $\mu$M. Equal volume of DMSO was used as a control.

Growth curves for each treatment were compared by two-way ANOVA.

*Flow cytometry*

Cell analyses and sorts were performed as described [8] (FACSCalibur, FACSCanto II, FACSAria III flow cytometers/cell sorters BD Biosciences). Subtraction of cell autofluorescence [13] and compensation of Alexa488-stained cells in the red channel was performed as described [14].

***Microscopy***

*Staining of living cells*

Living cells were stained with 1 µM of the lipid tracers DiOC, DiA, and DiLC. DiA induced rapid cell toxicity and was discarded from further analysis.

*Staining of fixed cells*

MTE4-14 cells plated on glass coverslips were fixed with 4% paraformaldehyde in PBS for 20 min. The paraformaldehyde was quenched with 50 mM $NH_4Cl$ for 20 min. The cells were then stained with DiOC, DiLC and an anti-CD9 Ab.

*Sample preparation for immunofluorescence*

The cells were seeded on glass coverslips and cultured for 24 h to 72 h, as indicated. For serum-starvation assays, the cells were seeded in complete medium. After 24 h, the cells were washed and refed with medium supplemented with 0.1% FCS, in which they were cultured/starved for 48 h. EGF, FGF-1, HGF, IGF-1, PDGF, SCF, or VEGF were added at 1, 10 or 100 nM, as indicated; after 30 min stimulation, the cells were fixed and stained. Cliff size, protein recruitment (CD9), and activating phosphorylation (p-PKCα) after starvation or growth factor addition were determined on co-stained slides by adapting an immunohistochemistry-proficient score-categorization procedure [15]. Cliff size and intensity of protein recruitment to cliffs were determined on 50 to 100 individual cells per group, using a five-class categorization of the cell perimeter fraction occupied by cliffs (from 1, nil/near nil, to 5, essentially the entire cell perimeter), and the staining intensity at cliffs (from 1, barely detectable, to 5, the highest intensity) (Figure 7, Table S3). Perimeter scores were multiplied by the corresponding intensity scores, and the square root of the product was taken, for a final five-class categorization of each cell. This procedure was performed by ≥2 independent observers (V.R., R.T., S.A.) who were blind to sample identity, on a cell-by-cell basis. Conflict in class value determination was solved by group discussion.

The cells were treated with 200-2000 nM cytochalasin D as described [10]. Cell treatment with 50 μM colchicine was performed essentially as described [16]. In parallel assays cells were treated with 10 μM blebbistatin, 200 nM latrunculin or 10 μM nocodazole. Phorbol 12-myristate 13-acetate ester was added at 200 nM for 30 min at 37 °C. Recovery from phorbol ester treatment was assessed after cell washing and post-incubation for 6 h or 24 h at 37 °C. Live cells were labeled with 4 μM calcein AM for 1 h at 37 °C.

*Confocal microscopy*

Slides were analyzed with one of two confocal microscopes (LSM510 META; LSM800). Three laser beams were used, emitting at wavelengths of 488 nm (argon ion laser, 200 mW, 2%-5% applied laser power), 543 nm (diode laser, 1 mW, 50%-100% applied laser power), 633 nm (diode laser, 5 mW, 50% applied laser power), except where indicated. HFT 488/543 or 488/543/633 beam-splitters were used, as needed for multi-color fluorochrome-conjugated antibody analysis, with band-pass emission filters 505 nm to 550 nm (green channel); long-pass 560 nm (for two-color analysis), or band-pass 560 nm to 615 nm (for three-color analyses) in the orange channel; long-pass 650 nm in the deep-red channel. HFT 488/543, and a band-pass of 505 nm to 550 nm filter in the green channel were used for EGFP/YFP, together with a long-pass at 560 nm for mCherry detection. Images were acquired in Multiplex mode, i.e., via sequential acquisition of individual laser lines/fluorescence channels, rather than by simultaneous acquisition of signals of distinct fluorophores, as simultaneously excited by different laser lines, to prevent cross-channel fluorescence spill-over. Detector gains of ≤770 V were applied to minimize electronic noise. Amplifier gains were ≤2.8×. Three-dimensional reconstruction was performed with the ZEN 2009 Light Edition software.

*Fixed cell analysis*

Cells cultured on glass coverslips were fixed with 4% paraformaldehyde in PBS for 20 min at room temperature. Permeabilization and blocking were performed in medium with 10% FCS and 0.1% saponin [17]. For staining of target protein extracellular domains, live cells on glass coverslips were stained in medium with 10% FCS at 37 °C for 5 min, and fixed after staining with paraformaldehyde, for 20 min at room temperature, or methanol-acetic acid (3:1), twice for 6 min at -20 °C. Cell membranes stained as above with lipophilic tracers were analyzed for triple fluorescence with excitation of DiOC at 488 nm, of DiLC at 543 nm, and for anti-CD9-Alexa633 at 633 nm.

For β-actin staining, cells were incubated with phalloidin-FITC or phalloidin-TRITC in staining medium containing 0.1% saponin at 10 μg/mL for 30 min at room temperature. Alternatively, cells were transfected with Lifeact, as composed of the first 17 amino acid from the *S. cerevisiae* Abp140, an actin-binding protein conjugated with mRFP1 (Ibidi GMBH). Lifeact stains filamentous actin (F-actin) in eukaryotic cells, without interfering with actin dynamics [18].

All images were acquired as 1024 × 1024 pixel format, except where indicated. Images were captured as averages of four sequential acquisitions, using a 63×/1.4 oil DIC objective (Plan-Apochromat; Zeiss). Nine to 14 contiguous cliff segments were analyzed in images of cells stained with lipophilic tracers and an anti-CD9 mAb, which were acquired as 2048 × 2048 pixel format (Table S1). Cell details were analyzed by applying a 1.7-3.7× zoom. Z-stacks were acquired as 512 × 512 pixel format, 0.5-0.8 $\mu$m interval between slices. The 512 × 512 pixel size was from 120 nm to 280 nm, the voxel size of corresponding Z-stacks was 0.120×0.120×0.800 $\mu m^3$ to 0.280×0.280×1.000 $\mu m^3$ (X,Y,Z). The pixel size of 1024 × 1024 pixel format was from 60 nm to 140 nm, that of 2048 × 2048 pixel frames was from 30 nm to 70 nm.

*Protein/lipid ratios*

Cliff tracers were quantified in representative cells transfected with CD9 and stained with DiOC and DiIC, and anti-CD9 antibody–Alexa633. Segmentation of adjacent high versus low signal intensity areas that delimited the cell contour were drawn as described in Table S1. The mean intensity of DiOC, DiIC, and CD9 signals was computed for each cliff segment, after conversion to a normalized gray scale and average background subtraction. Signal volume (surface area × gray signal intensity) was computed and converted to percentage intensity, i.e. the fraction of total cliff signal associated to the segment analyzed within each individual cell. The average of the DiOC and DiIC signals in each membrane region was normalized to 1. The ratio of the CD9–mCherry signals versus DiOC + DiIC in the corresponding regions was then computed.

*Live cell analysis*

Live cells cultured on glass slides were analyzed in Leibovitz's F15 culture medium without phenol red and bicarbonate, supplemented with 10% FCS, 100 IU/mL penicillin, 100 $\mu$g/mL streptomycin, 25 mM HEPES

and 2 mM N-acetyl-cysteine, to reduce free-radical damage and pH increase. Excitation of CFP was at 458 nm, that for EGFP/YFP was at 488 nm, that for mRFP1 and mCherry was at 543 nm. Images were captured at 30 s/1 min intervals, except where indicated, using a 63×/1.4 oil DIC objective (Plan-Apochromat; Zeiss).
The amount of proteins or protein–fluorescent protein chimeras at distinct cell/cell membrane sites was quantified using the ImageJ/Fiji software on the acquired frames. Reference curves for signal quantification were power curves fitting optical density gray-scale standards by Kodak.
The calcium-calmodulin-binding peptide (designated as genetically encoded $Ca^{2+}$ indicators for optical imaging; GECO) was composed of a myosin light-chain kinase fragment that binds calmodulin (M13), a circularly permuted fluorescent protein and vertebrate calmodulin. G-GECO1.2 included GFP, R-GECO1.2 included mRFP1. The plasmids CMV-G-GECO1.2 (Addgene, #32446) and CMV-R-GECO1.2 (Addgene, #45494) were kindly provided by R. Campbell. The plasmid Lifeact–mRFP1 was a kind gift from G. Scita. Signaling by PKCα-GFP, PKCδ-GFP, PKCε-GFP was induced with the 162-46.2 anti-Trop-2 mAb Signaling path integrity/PKC stimulation was conducted with ATP 5 $\mu$M [19].

*Antibody-mediated co-capping*

Cells were detached from culture plates using trypsin, and incubated with primary antibodies against Trop-2 for 20 min on ice. After washing, cells were incubated with the secondary Alexa Fluor 488-conjugated antibodies for 10 min at 37 °C, to cross-link target molecules and to induce capping of the antigen–antibody complex. Co-clustering with other proteins under study was expected in case of tight interactions with the molecules undergoing capping [20]. The 'capped' cells were fixed with 1% paraformaldehyde for 10 min at room temperature. The paraformaldehyde was quenched with FCS, and the cells were stained with antibodies against the membrane protein(s) of interest. The co-capping of resident molecules typically appeared as a yes-or-no phenomenon, and was listed as occurrence versus non-occurrence.

*Single-molecule near-field optical microscopy (NSOM) analysis*

NSOM imaging experiments were performed as described [21]. Human MCF-7 cells were seeded on fibronectin-coated glass-bottomed 35 mm μ-Dish High Grade (Ibidi, Martinsried, Germany). After 24 h, the culture medium was replaced with phenol-red-free complete medium supplemented with 25 mM HEPES buffer for optimal image acquisition and stained with HT29/26-Alexa 633 anti-Trop-1 ($\lambda_{ex}$= 568nm, $\lambda_{em}$> 620nm, scan speed 2 ms/pixel) and T16-Alexa 546 anti-Trop-2 mAb ($\lambda_{ex}$= 530nm, $\lambda_{em}$> 560nm, scan speed 1ms/pixel).

### Statistics

Student's t-tests were used for comparisons of mean platform length across (colocalized) molecule cliffs. Two-tailed Fisher exact tests were used to compare protein expression levels in nontransfected versus gene-transfected cancer cells. Spearman nonparametric correlation coefficients were computed. Data description for categorical variables p-PKCα and CD9 is by bar plots and standard deviation bars at each time of measurement.

Comparisons between the distributions of the variables at time 0 (control cells) versus treatment time and recovery was performed by $\chi$-squared statistical analysis. Data descriptions of the paired differences between variables P-PKC$\alpha$ and CD9 according to times of measurement were performed using asymmetric confidence intervals, over proportions based on logit transformation. Data were analyzed using Sigma Stat (SPSS Science Software UK Ltd.), GraphPad Prism (GraphPad Software Inc., USA), and SISA (www.quantitativeskills.com/sisa/statistics/).

**Supplemental Figures**

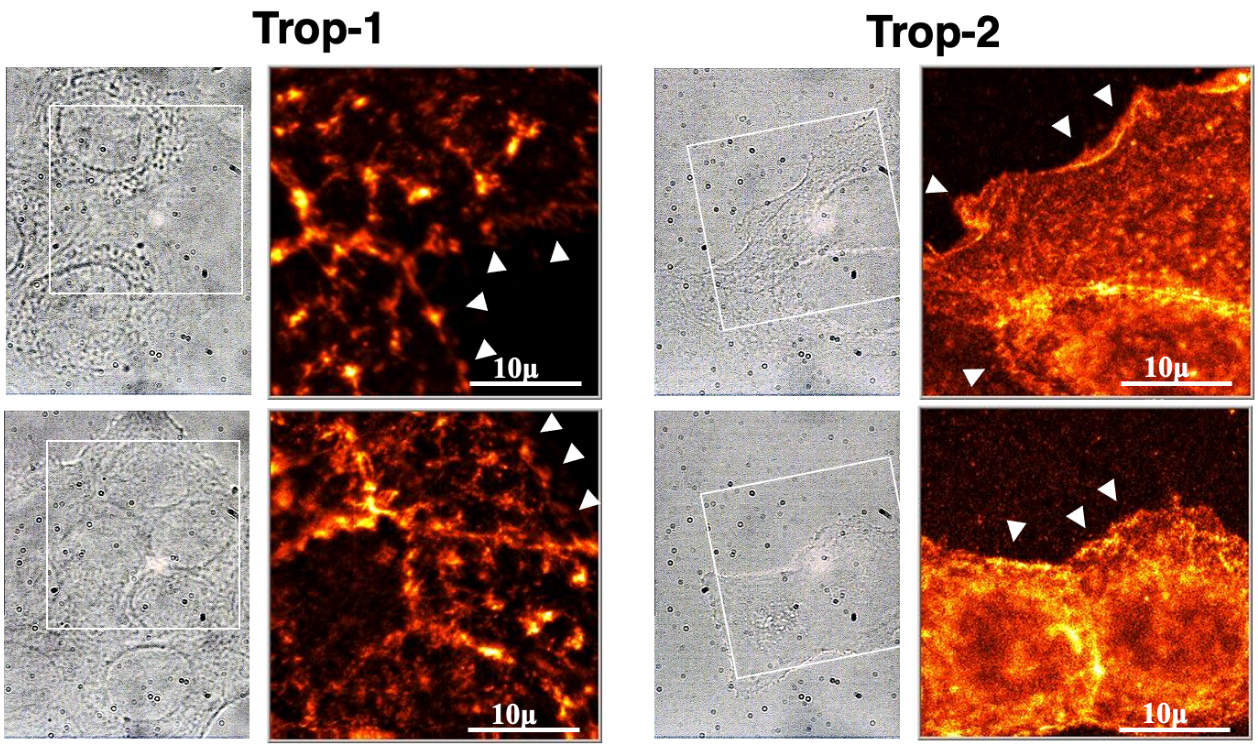


**Figure S1. NSOM analysis of Trop-1 and Trop-2 at cliffs.** (Related to Figure 1) Images were acquired as described in Material and methods. Trop-1: $I_{ex}$~ 1.3kW/cm$^2$, $I_{max}$= 5000 top image, $I_{ex}$~ 1.0kW/cm$^2$, $I_{max}$= 400 bottom image. Trop-2: $I_{ex}$~ 150W/cm$^2$, $I_{max}$= 150 top image, $I_{ex}$~ 150W/cm$^2$, $I_{max}$= 80 bottom image. Images showed a localization accuracy of approximately 3 nm. (*left images*) ROI are indicated by white squares. (*right images*) Pseudo-color representation of Trop expression levels. Yellow: highest density. Black: no signal. White arrowheads: cliffs. Bars, 10 µm.

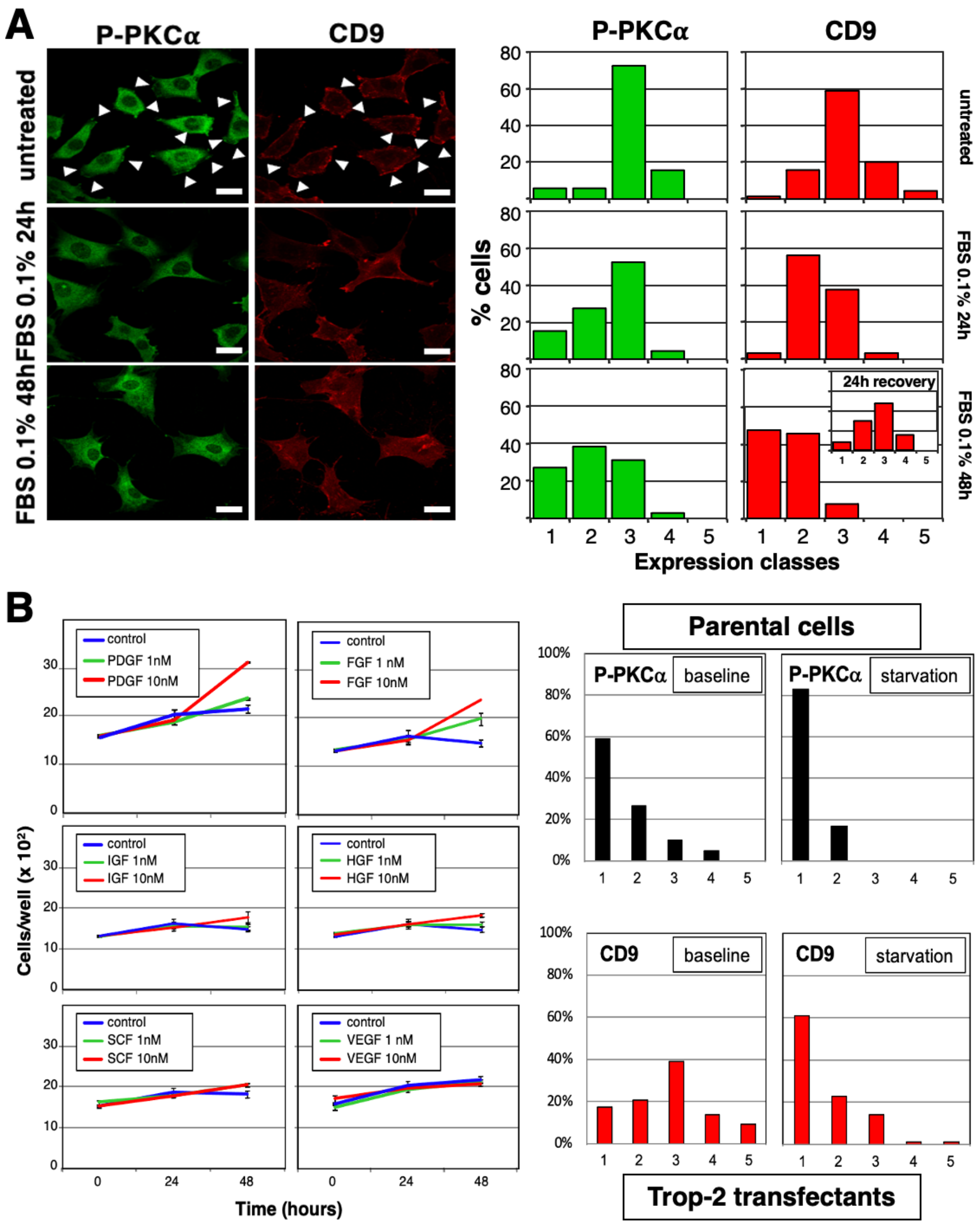


**Figure S2. Growth factors induce membrane cliffs.** (Related to Figure 7, Table S3). White arrowheads indicate areas of colocalization of signal transducers. (**A**) (*left*) MTE4-14 cells were growth factor-starved (0.1% serum) for 24 h or 48 h and co-recruitment of P-PKCα and CD9 at cliff sites was quantified by image analysis of cells stained with anti-P-PKCα Alexa488 and anti-CD9

Alexa633 mAb. Cliffs were found to disappear after 24 h of serum starvation. Bars, 20 µm. (*right*) Cultured cells stained with anti-P-PKCα and anti-CD9 mAb were classified into 5 expression classes, as described. Progressive disappearance of cliffs over time upon starvation was observed. Addition of 10% serum to serum-deprived cells led to full recovery of cliffs in 24 h (*bottom right,* inset). The full datasets and comparison statistics are presented in Table S3. (**B**) (*left*) Serum-starved MTE4-14/Trop-2 cells were treated with GF at the concentrations indicated. PDGF and FGF-1 induced cliffs and cell growth. HGF, IGF-1 showed lesser impact on both cliffs and cell growth, SCF, VEGF had no effect on either parameter. (*right*) Expression of Trop-2 was shown to induce endogenous CD9 at baseline. Loss of both P-PKCα and CD9 signals at cliffs was observed upon cell starvation.

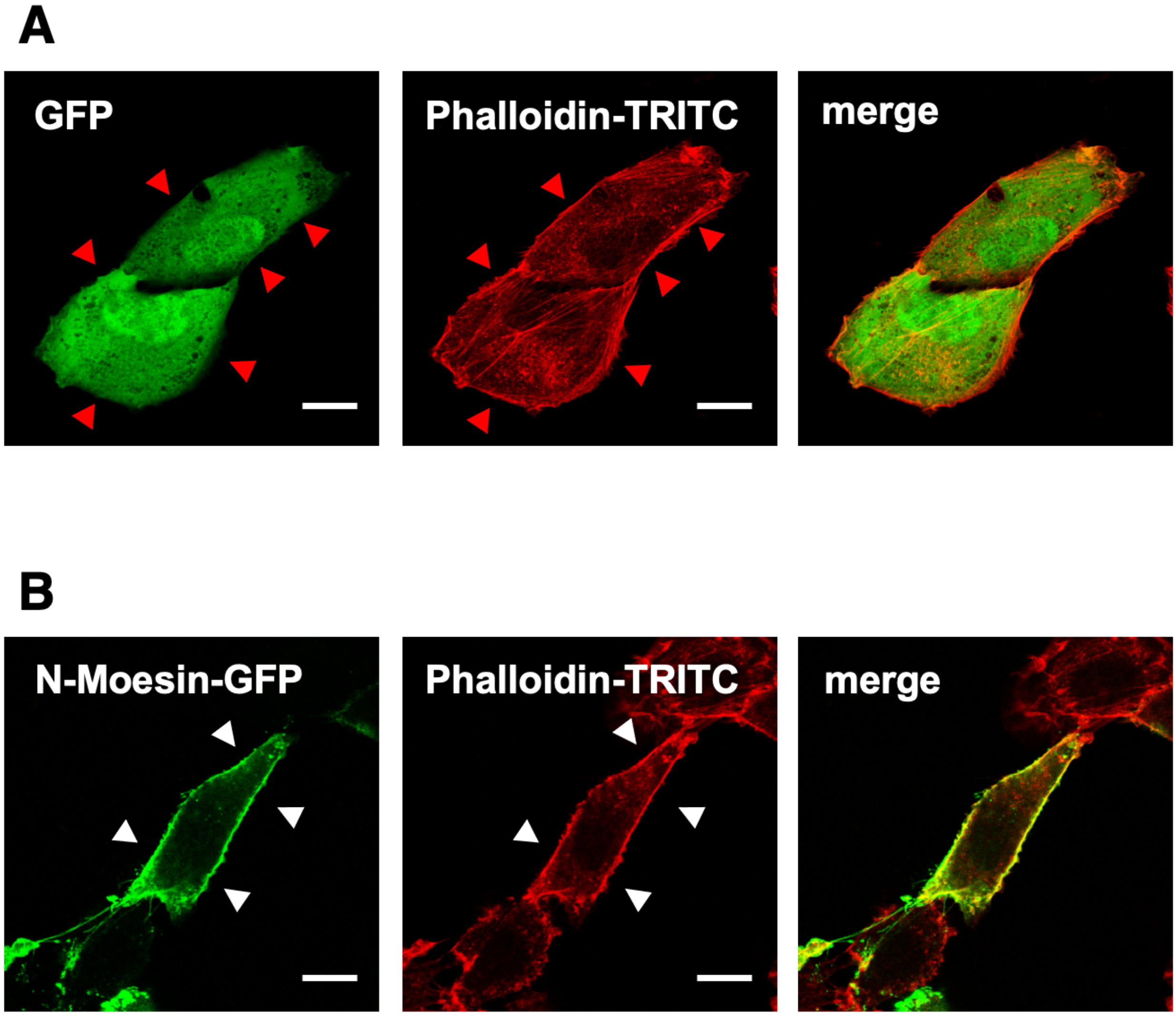


**Figure S3. The β-actin cytoskeleton scaffolds membrane cliffs.** (Related to Figures 1, 2). (**A**) MTE4-14 cells were transfected with GFP as a cytoplasmic tracer and as a negative control for recruitment at cliffs. β-actin was labeled with falloidin-TRITC. (**B**) MTE4-14 cells transfected with N-moesin-EGFP. β-actin was labeled with falloidin-TRITC. Merged signals are indicated on the right panels. Colocalization at membrane cliffs is indicated by white arrowheads. Absence of colocalization at cliffs is indicated by red arrowheads. Bars, 5 µm.

### Movies

#### Movie S1. Co-recruitment of CD9 and Trop-2 at cliffs

(Related to Figures 1, 2) MTE4-14 cells transfected with Trop-2–GFP (green) and CD9–mCherry (red) were imaged over time. Acquisition time for each image was 2.5 s. The time-lapse covers a period of 5 min. Sites of recursive colocalization Trop-2–GFP and CD9–mCherry at cell membrane cliffs are indicated by arrowheads.

#### Movie S2. Recursive co-recruitment of Trop-2 and ERK at cliffs

(Related to Figures 1, 2) MTE4-14 cells transfected with ERK–GFP (green) and Trop-2–mRFP1 (red) were imaged over time. Acquisition time for each image was 2.5 s. The time-lapse covers a period of 5 min. Sites of recursive colocalization at cell membrane cliffs are indicated by arrowheads.

#### Movie S3. 3D reconstruction of cell membrane cliffs

(Related to Figure 3, Movie S2) Three-dimensional reconstruction of an MTE4-14 cell transfected with wtTrop-2 and stained with the T16 anti-Trop-2 mAb. A living cell was stained, to allow exclusive visualization of outer cell-membrane Trop-2. The live cell was fixed after staining with 4% paraformaldehyde in phosphate-buffered saline for 20 min. A Z-stack of nine 1024×1024 pixel format images (pixel size 100 nm; voxel size 0.100×0.100×0.500 $\mu m^3$; X,Y,Z) was reconstructed in a three-dimensional rendering with the Volocity software. Cliffs are the highest signal-density areas. Membrane cliffs are the elevated membrane regions detected at cell sides. The three largest cliffs are enclosed in white ovals (details are presented in Figure 3).

#### Movie S4. Dock appearance and space-time dimensions are unrelated to cell movement

(Related to Table S2) MTE4-14 cell transfected with CD9–mCherry were imaged by time-lapse confocal microscopy. The recruitment of CD9-mCherry at docks is indicated by white arrowheads. Images were taken at intervals of 4.8 min. The acquisition time for each image was 2.53 s. The time-lapse covers a period of 10 hours. No correlation was detected between dock appearance parameters (size, location, periodicity) and extent or direction of cell movement.

#### Movie S5. Four-dimensional cliff dynamics

(Related to Figure 1, Movie S3) Four-dimensional rendering (three-dimensional, 3D reconstructions over time) of an MTE4-14 cell transfected with CD9–mCherry. Each Z-stack is composed of nine images, 512×512 pixel format (pixel size 140 nm; voxel size 0.140×0.140×0.800 $\mu m^3$; X,Y,Z); acquisition time for each image was 1.28 s; images were taken every 26.6 s. The time lapse covers a period of about 35 min. Z-stacks were reconstructed as a sequence of 3D images, generated with the Zen software. The three largest cliffs are enclosed in white rectangles. A pseudo-color (LUT from Fiji/ImageJ) representation of the MTE4-14 cells was used to highlight the variation of absolute amounts of CD9 at cliffs (deep blue corresponds to the highest levels).

**Movie S6. Signal triggering induces $Ca^{2+}$ waves from cliffs**

(Related to Figures 1, 3) MDA-MB-231 cells were transfected with the $Ca^{2+}$ probe G-GECO and analyzed by time-lapse confocal microscopy under resting conditions and after signal induction with the 162-46.2 anti-Trop-2 mAb. A $Ca^{2+}$ wave originated at membrane cliffs (white arrowheads) and diffused directionally throughout the cell cytoplasm. Acquisition time for each image was 2.53 s. Images were taken in continuous mode.

**Movie S7. PKCα signaling is induced at cliffs**

(Related to Figures 1, 2) OvCa-432 ovarian cancer cells were transfected with PKCα-GFP (green) [22] and analyzed by time-lapse confocal microscopy under resting conditions and after signal induction with the 162-46.2 anti-Trop-2 mAb. PKCα-GFP recruitment at cliff segments was recorded over time. The time-lapse covers a period of about 5 min. Acquisition time for each image was 2.5 s.

**Movie S8. Kinase-inactive PKCα recruitment at cliffs**

(Related to Figures 1, 2) MTE4-14 cells were transfected with dominant-negative K368R PKCα-GFP (green) [22] and CD9-mCherry (red) and imaged by time-lapse confocal microscopy. K368R PKCα-GFP recruitment at CD9-mCherry cliffs is indicated by white arrowheads. Acquisition time for each image was 2.5 s. The time-lapse covers a period of about 5 min.

**Movie S9. Co-recruitment of the β-actin cytoskeleton and Trop-2 at cliffs**

(Related to Figure 8) MTE4-14 cells transfected with Trop-2–GFP (green) and the β-actin cytoskeleton marker Lifeact–mRFP1 (red) were imaged over time. (*right*) Merged images show the overlap of green and red fluorescence at cliffs. Acquisition time for each image was 2.5 s. The time-lapse covers a period of about 5 min.

**Movie S10. Assembly of the β-actin cytoskeleton at cliffs parallels induction of Ca2+ signaling**

(Related to Figure 8) OvCa-432 ovarian cancer cells were transfected with the β-actin cytoskeleton marker Lifeact–EGFP (green) and with the $Ca^{2+}$ probe R-GECO (red) and analyzed by time-lapse confocal microscopy under resting conditions and after signal induction with the 162-46.2 anti-Trop-2 mAb. Acquisition time for each image was 2.53 s.

**Movie S11. Integrity of the β-actin cytoskeleton is required for membrane cliff scaffolding**

(Related to Figure 8) MTE4-14 cells transfected with PKCα-GFP and CD9-mCherry were treated with 200 nM cytochalasin D and analyzed by time-lapse confocal microscopy. Acquisition time for each image was 2.53 s. Images were taken in a continuous mode. Progressive disappearance of PKCα-GFP and CD9-mCherry membrane cliff signal was observed during the cytochalasin D treatment.

**Movie S12. Microtubule integrity is not required for membrane cliff scaffolding**

(Related to Figure 8) MTE4-14 cells transfected with PKCα-GFP and CD9-mCherry were treated with 50 μM nocodazole and analyzed by time-lapse confocal microscopy. Acquisition time for each image was 2.53 s. Images were taken in a continuous mode. Essentially unvaried PKCα-GFP and CD9-mCherry membrane cliff signal was observed during the nocodazole treatment.
